# Integrative metabolomics reveal the organisation of alkaloid biosynthesis in *Daphniphyllum macropodum*

**DOI:** 10.1101/2022.05.25.493403

**Authors:** Kaouthar Eljounaidi, Barbara Radzikowska, Caragh Whitehead, Susana Conde, William Davis, Adam Dowle, Swen Langer, Tony Larson, William P. Unsworth, Daphne Ezer, Benjamin R. Lichman

## Abstract

*Daphniphyllum* alkaloids are structurally diverse nitrogen-containing compounds with polycyclic, stereochemically rich carbon skeletons. Understanding how plants biosynthesise these compounds may lead to greater access to allow exploration of bioactivities; however, very little is known about their biosynthetic origins. Here, we integrated metabolomics approaches to map alkaloid distribution across *Daphniphyllum macropodum* plants and tissues. We generated a novel untargeted metabolomics workflow to highlight trends in alkaloid distribution across tissues, using a holistic approach that does not rely on ambiguous peak annotations. Both liquid-chromatography-mass spectrometry and mass-spectrometry imaging analyses independently revealed that alkaloids have a pattern of spatial distribution based on their skeletal subtypes. The distinct alkaloid subtype localisation suggests the biosynthetic pathway is controlled spatially with intermediates transported from the phloem to the epidermis where they undergo additional derivatization. This study sets the stage for the future work on *Daphniphyllum* alkaloid biosynthesis and highlights how integrating different metabolomics strategies can reveal valuable insights on these compounds’ distribution within the plant.

## Introduction

To adapt to their environment, plants have evolved to biosynthesise a plethora of small organic molecules, often referred to as natural products or specialized metabolites. Among these compounds, many alkaloids (nitrogen containing natural products) have profound biological activities and are widely used as therapeutics and analgesics.

Plants from the genus *Daphniphyllum* are dioecious shrubs or small trees native to tropical and subtropical Asia. These evergreens are often grown as ornamental plants due to their characteristic red-plum petioles and dark green leaves (*1*). Some *Daphniphyllum* species have also been used in Chinese traditional medicines to treat asthma, rheumatism, and snake-bites (*2*). *Daphniphyllum* sp. are known for their ability to produce structurally diverse and complex alkaloids characterised by polycyclic cage-like structures, and stereochemically rich molecular frameworks (Fig. 1A) (*3*). Whilst several alkaloids have demonstrated bioactivities including anti-inflammatory (*4*), neuroprotective (*5*) and anticancer (*6–8*) they are still significantly understudied in spite of their enormous structural diversity.

**Figure 1.**
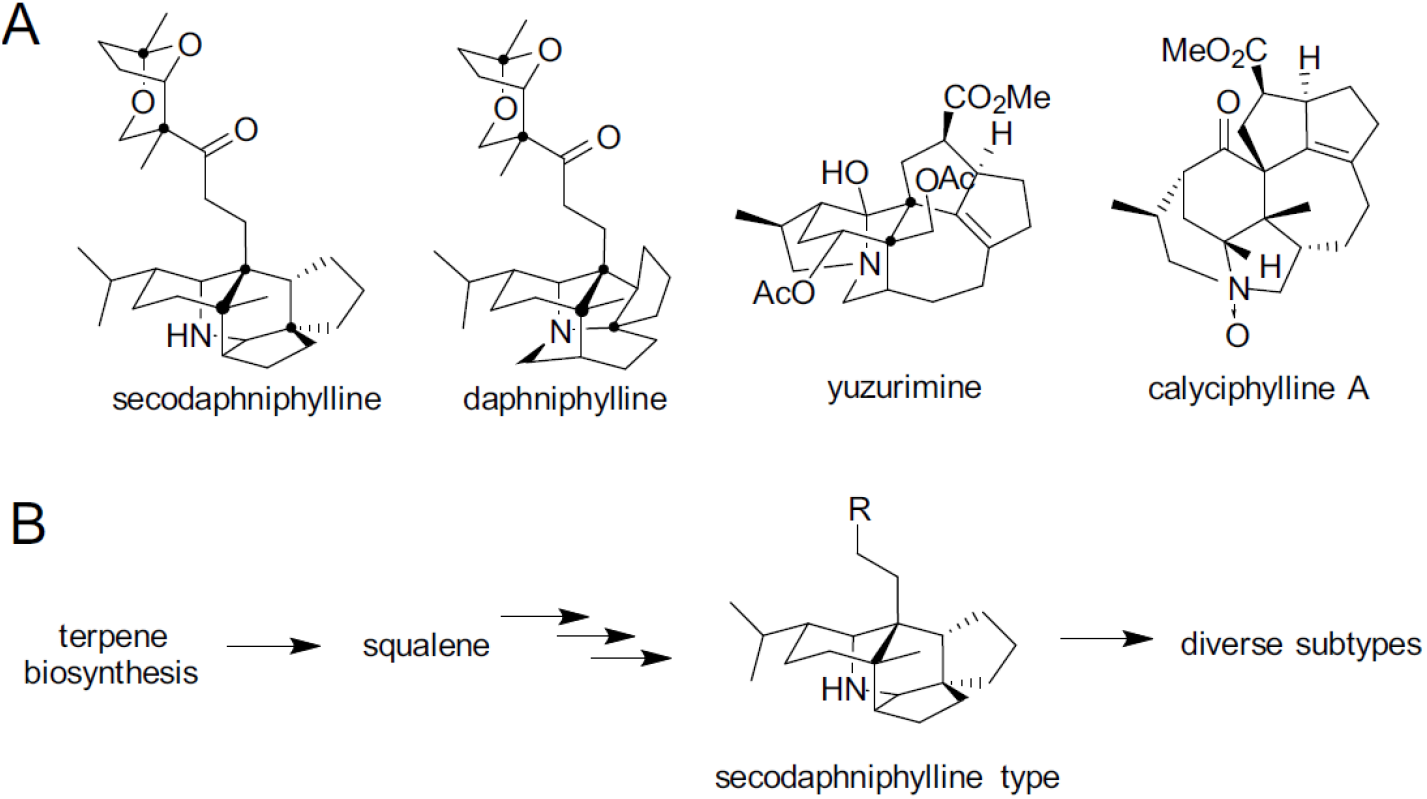
*Daphniphyllum* alkaloids. A. Representative *Daphniphyllum* alkaloids found in *D. macropodum*. B. Proposed pathway via squalene and a secodaphniphylline type precursor.

More than 330 alkaloids have been identified in these plants to date. Based on their skeletal structures, they can be categorised into 13-35 structurally distinct type subclasses including daphniphylline, secodaphniphylline, calyciphylline-A, yuzurimine, and yuzurine types (*9*). Due to their structural complexity and potential biological activities, *Daphniphyllum* alkaloids (DAs) have remained targets for total synthesis efforts over many decades (*10–15*).

Feeding experiments (*16*) and pioneering biomimetic syntheses by Heathcock (*10*) suggested that DAs are terpenoids, deriving from the mevalonic acid pathway and specifically from squalene. Triterpene alkaloids are rare, with most examples being steroidal alkaloids such as solanine from Solanaceae and cyclopamine from *Veratrum californicum* (*17–19*). Biomimetic conversions indicates that secodaphniphylline-type compounds are precursors to all other DAs (Fig. 1B) (*20, 21*).

Increasing knowledge about these compounds and how they are made in the plant will facilitate bioengineering strategies to increase their availability for further bioactivity screens. New insights about their biosynthesis will also provide access to novel enzymes and biocatalysts that can be used in the assembly of complex stereochemically rich structures.

Here, we integrated metabolomics approaches, namely high-resolution liquid-chromatography tandem mass spectrometry (HR-LC-MS/MS) and MS-imaging, to map alkaloid distribution within *Daphniphyllum macropodum*. This comprehensive analysis provides insights into the spatial allocation of biosynthetic pathways, which can be used to understand and eventually reconstitute *Daphniphyllum* alkaloid biosynthesis. We developed a novel workflow for untargeted metabolomics to leverage DA *in silico* databases in the absence of chemical standards. Our analysis revealed alkaloids are distributed across plants and tissues based on their skeletal subtypes, with yuzurimine subtypes accumulating primarily in younger leaves and in the epidermal cells, whilst secodaphniphylline and daphniphylline subtypes appear to be enriched in plant material containing vascular tissue, and correspondingly are found to localise to tissue in the phloem region. This led to the development of a biosynthetic model where secodaphniphylline subtype compounds are initially located in phloem tissues, then transported to the epidermis where they undergo additional synthetic steps to yield yuzurimine type compounds. This study sets the stage for the future work on *Daphniphyllum* alkaloids biosynthesis and highlights how integrating different metabolomics strategies can reveal valuable insights on compounds’ distribution within the plant.

## Methods

### Database construction

An *in silico Daphniphyllum* natural product database was manually assembled, reporting chemical names, synonyms, molecular formula and structural formula. Entries were obtained searching Reaxys (https://www.reaxys.com/, Elsevier) for substances “Isolated from Natural Source: Daphniphyllum”. We also gathered entries from two recent comprehensive reviews and research articles (*3, 21*). Dereplication and correcting ambiguous entries was performed manually. Each *Daphniphyllum* alkaloid was assigned to a structural subtype using definitions from Chattopadhyay and Hanessian (*3*).

### Plant and Tissue selection

Plant material was obtained from a variety of sources. Plants A-D are located in Ray Wood on the grounds of Castle Howard, North Yorkshire, UK, managed by the Yorkshire Arboretum. Plant A (accession 197831591) is a male *Daphniphyllum macropodum* originally from a horticultural source (Treseder Nurseries) planted in 1978. Plant B (accession 19879448) is a female *Daphniphyllum macropodum* var *humile* from Japan planted in 1989. Plants C and D, female and male respectively, were planted in 1989 and are from Fanjing Shan area of the Guizhou province, China. They were verified as *Daphniphyllum macropodum* by Bleddyn Wynn-Jones of Crûg Farm Plants, Wales, UK (personal communication). Plants E and F, both *Daphniphyllum macropodum*, were purchased from Burncoose Nurseries, UK, and Architectural Plants, UK, respectively and grown in the University of York. They were planted in 2018 and both originate from seed taken from mature specimens in the UK. Plant G, also grown at the University of York, is *Daphniphyllum macropodum* var *humile* (BSWJ11232) purchased from Crûg Farm Plants, Wales, UK. Our specimen was planted in 2018; the progenitors originate from Japan, collected in 2005. Plant tissue was obtained from plants through 2019 and 2020. Tissues were sampled onto dry ice before storing at −70 °C.

### LC-MS extraction method

Plant material was freeze dried and ground with pestle and mortar. Each tissue was sampled in triplicate. The sample (~10 mg) was extracted with 80% methanol (500 μL) and vortexed thoroughly. The samples were then shaken in a TissueLyser (Qiagen) (10 min, 5 Hz) and then centrifuged (4000 x g, 2 min). Supernatant (50 μL) was diluted 10x with 80% methanol (450 μL) and added to a 96 well plate. The plates were sealed and stored at 4°C.

### LC-MS run method

LC-MS/MS runs were performed using a Waters Acquity I-Class UPLC instrument interfaced to a Thermo Tribrid Fusion Orbitrap instrument using an APCI ion source in positive ion mode. UPLC runs were controlled by Waters Empower software. 2 μL injections were made onto an Acquity BEH C18 column (100 x 2.1 mm, 1.7 um particle size), held at 60 °C. The column was run at 0.5 mL/min with mobile phase A = 10 mM ammonium bicarbonate (pH 10.2) and B = MeOH, under the following gradient program: initial isocratic 2% B; 0.2-0.5 min linear to 40% B; 0.5–4 min linear to 80% B; 4–4.5 min isocratic 80% B; 4.5–4.6 min return to initial conditions and hold until 5 min (total run time). MS parameters were controlled by Thermo Xcalibur 4.1 software. Eluent flow was diverted into the APCI source between 0.66 and 4.5 min. The source was maintained at 350 °C and a spray current setting of 4 uA. Nitrogen was used as sheath, aux, and sweep gas, set to 35, 5, and 0.5 arbitrary units, respectively. The ion transfer tube was held at 275 °C. Data was collected in data-dependent MS^2^ mode. MS^1^ data was collected in profile mode using a cycle time of 0.4 s and a desired minimum points across the peak setting of 6, over a *m/z* range of 200-2000 with orbitrap resolution set to 60 000 (FWHM @ 200 *m/z*). Easy-IC internal calibration was used to reduce MS^1^ mass errors to ~< 1 ppm. Low resolution data-dependent MS^2^ data was collected as centroided data in rapid scan mode in the ion trap, using alternating CID and HCD fragmentation modes and precursor quadrupole isolation windows set to 1 *m/z*. HCD data was collected at stepped normalised collision energies of 34, 40, and 45%, and CID data was collected using a fixed normalised collision energy of 40%. Dynamic exclusion was set to collect one CID and HCD MS^2^ scan from each precursor ion and to exclude further fragment scans from the same precursor for 2 s.

### Processing LC-MS peaks

Files were converted from .raw to .mzML format using the Proteowizard msConvert tool, and subsequently processed using bespoke scripts in R 4.0.3 in a linux environment. Briefly, the R package XCMS (3.12.0) was used to detect features using the “centWaveWithPredictedIsotopeROIs” method, with the following settings: ppm = 10; snthresh = 10; peakwidth = 2, 10; prefilter = 3, 1000; integrate = 2; mzdiff = −0.1. Only features collected between 0.2 and 4.5 min were retained, and feature detection errors (duplicates, poor gaussian peak shapes) were identified and those features removed using custom scripts. Features between samples were aligned and grouped across samples using modified versions of the XCMS obiwarp() and group() functions, respectively. Grouping required that all features were detected within replicate group. Missing features were imputed by re-integrating the raw data using a modified version of the fillPeaks() function. The feature list was dereplicated using the CAMERA package, with all samples except blanks used as inputs.

CAMERA (1.33.3) (*22*) was run in sequential steps using the following CAMERA functions as follows: groupFWHM() with perfwhm set to 0.5; modified findIsotopes() with maxcharge = 2.maxiso = 4, ppm = 5; groupCorr() using graphMethod = “lpc” and calcCiS only (feature shape analysis); groupCorr() with graphMethod = “lpc” and calcCaS only (correlation between samples). Correlation thresholds for calcCiS and calcCaS steps were set to 0.75. Finally, findAdducts() was used with the default settings. Candidate formulae for detected ions were generated using the rcdk (3.5.0) package, using the following elemental limits: C 2-42; H 0-62, O 0-18; N 0-3; Na 0-1. Formulae were further constrained to be with the RDBE range of 0.5, 18 and within 5 ppm mass error.

The final feature set was filtered as follows: 1) A cutoff area value was calculated for each feature, defined as the mean plus 3 x 1 SD of the area across blank runs. A feature in at least one non-blank sample had to exceed this cutoff to be retained; 2) Any filtered feature that has a MS^1^ *m/z* match within 5 ppm to the in-house database is retained as a potential hit. This database included 404 compounds identified from the literature (mostly alkaloids with some phenylpropanoids), with exact masses calculated for [M+H]^+^, [M+NH4]^+^, and [M+Na]^+^ ions; 3) For the remaining features, where multiple isotopes and adducts were detected by CAMERA across a set of grouped features, only a single representative feature with a valid formula was kept (usually most intense likely monoisotopic ion).

Areas from the MS^1^ feature list were then processed by subtracting the blank cutoff values and normalising to dry weight. Finally, these values were loess normalised to the QC samples using the R MetMSLine (1.2.1) package. As the QCs were not pooled samples, but representative samples, not all features over the data set were present in the QC samples. Per-feature based loess normalisation was therefore not possible with this dataset without imputing missing values into the QCs. We proceeded as follows: 1) For QC samples, replace any feature where > 80% of the values are zero with the mean value; 2) calculate the loess fit across the QCs and samples using 7-fold cross-validation; 3) Replace any computed negative values with zero. The performance of the normalisation was evaluated by examining improvements in SD values across replicates.

For subsequent analysis of MS^2^ data in GNPS, CID and HCD spectra associated with the filtered MS1 feature set were collated into .mgf files using custom R scripts. CID and HCD MS^2^ spectra were analysed separately in GNPS. Additionally, .mgf files were created using either composite MS^2^ spectra (from multiple files identified as belonging to one MS^1^ feature), or the best representative MS^2^ spectrum from a single file (the MS^2^ spectrum from a single file that is closest to the MS^1^ feature apex).

### Molecular Networking and Spectral Library Search

A molecular network was created with the Feature-Based Molecular Networking (FBMN) workflow (*23*) on GNPS (https://gnps.ucsd.edu, (*24*)). The mass spectrometry data were first processed with XCMS3 (*22, 25*) and the results were exported to GNPS for FBMN analysis. The data was filtered by removing all MS/MS fragment ions within +/- 17 Da of the precursor *m/z*. MS/MS spectra were window filtered by choosing only the top 6 fragment ions in the +/- 50 Da window throughout the spectrum. The minimum fragment ion intensity in the MS/MS spectra was set to 100. The precursor ion mass tolerance was set to 0.02 Da and the MS/MS fragment ion tolerance to 0.4 Da. A molecular network was then created where edges were filtered to have a cosine score above 0.7 and more than 6 matched peaks. Further, edges between two nodes were kept in the network if and only if each of the nodes appeared in each other’s respective top 10 most similar nodes. Finally, the maximum size of a molecular family was set to 0, and the lowest scoring edges were removed from molecular families until the molecular family size was below this threshold. The spectra in the network were then searched against GNPS spectral libraries (*24, 26*). The library spectra were filtered in the same manner as the input data. All matches kept between network spectra and library spectra were required to have a score above 0.7 and at least 6 matched peaks. The molecular networks were visualized using Cytoscape software (*27*).

### Data Deposition and Job Accessibility

Upon peer-reviewed publication, the mass spectrometry data will be deposited on public repository (provide the deposition accession number), such as MassIVE or MetaboLights. The molecular networking job can be publicly accessed at https://gnps.ucsd.edu/ProteoSAFe/status.jsp?task=a8b1a6fc87684bf99deeae350513ea7c.

### LC-MS data analysis

The loess corrected peak areas were filtered by removing values where peaks were only detected in one of three technical repeats. Next, all peak values for peaks whose maximum value fell below the median of all peak maxima were removed. The average and log-transformed LC-MS peak areas were calculated from technical repeats:

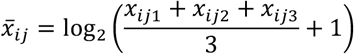

Where: *i* = sample, *j* = peak, *x* = raw peak area.

Then the peaks areas were normalised across all samples by dividing the averaged peak areas by root-mean-square value across all samples for a given peak.

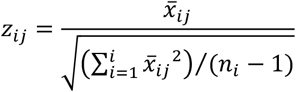

Where *z* = normalised peak area for *i* sample and *j* peak.

To describe the changes of a group of peaks in a given sample, the normalised peak areas, of peaks in that group, within a sample were summed.

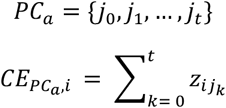

Where *PC_a_* (peak cluster or other collection of peaks) is a set containing *n_t_* of *j* peaks, *CE_PCa,i_* is compound enrichment of a given PC_a_ and sample *i. CE* is essentially a sum of the normalised peak values within a sample. Across multiple samples, an average of *CE* can be calculated. Due to the zero-inflated value of the original data, the transformed data is not normally distributed and non-parametric tests must be used for comparison of means (Kruskal-Wallis test, Dunn post-hoc test).

### Clustering algorithm

Averaged and log-transformed LC-MS peak areas (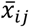 above) were used for determining cluster method, with the overall method structure being principle component analysis followed by UMAP followed by clustering. The following hyperparameters were tested using a grid search: UMAP components, UMAP neighbours, and number of Principal Components. Two different hierarchical clustering methods were used: complete or median. Consensus clustering with 20 starts was always used for K-means clustering. The clustering methods were compared using adjusted Rand-index (ARI) and methods with the highest ARI were selected. For sample clustering, the ground-truth *a priori* clusters were provided by plant and tissue type. For peak clustering, the ground-truth *a priori* clusters were based on chemical type annotation from MS^1^ and sub-network cluster index from the FBMN GNPS network based on HCD composite MS^2^ spectra.

To describe the subtype annotation within each peak cluster (PC) we first calculated a compound subtype score for each annotated peak based on the proportion of ambiguous annotations with particular subtypes. Each peak has a total score of one, but this is divided proportional to the annotation. For example, a peak annotated with three yuzurimine subtype compounds and one calciphylline A subtype compound would be scored 0.75 yuzurimine and 0.25 calciphylline A. To assign a subtype frequency across all peaks in a PC, these proportional subtype scores were summed.

### Purification of standards

^1^H and ^13^C NMR spectra were recorded on either a JEOL ECX400 or JEOL ECS400 spectrometer, operating at 400 MHz and 100 MHz respectively. All spectral data was acquired at 295 K. Chemical shifts (δ) are quoted in parts per million (ppm). The residual solvent peak, δ_H_ 7.26 ppm and δ_C_ 77.16 ppm for CDCl_3_ were used as a reference. Coupling constants (*J*) are reported in Hertz (Hz) to the nearest 0.5 Hz. The multiplicity abbreviations used are: s singlet, d doublet, t triplet, q quartet, quin. quintet, sex. sextet, m multiplet. Signal assignment was achieved by analysis of DEPT, COSY, HMBC and HSQC experiments where required. Purification of alkaloids was carried out using Interchim Puriflash 4250 prep HPLC system linked to an Advion Expression S Compact Mass Spec (CMS).

Plant leaf tissue (100 g) was ground to a fine powder under liquid nitrogen with a pestle and mortar. The powder was extracted, at room temperature whilst stirring, with methanol (350 mL for 2 h, then 150 mL for 16 h). The methanolic extract was filtered and dried *in vacuo*. The residue was resuspended in ethyl acetate (200 mL) which was extracted with aq. HCl (0.01 M, pH 2, 3 x 200 mL). The aqueous layers were combined and washed with ethyl acetate (200 mL). The aqueous layer was basified to pH 10 with sat. Na_2_CO_3_ and then extracted with chloroform (4 x 100 mL). The organic fractions were combined and dried *in vacuo* to yield crude alkaloid extract.

A Waters XBridge C18 5μm column of size 4.6 x 150 mm was used to analyse the extract using a gradient equilibrating column at 50-50% ammonium bicarbonate buffer pH = 10.2 and acetonitrile for 1 min, 50-50% to 5-95% over 6 min, holding 5-95% for 3 min with 1.4 mL/min flow. The peaks were detected using a MS detector. The samples were earlier prepared in the concentration of 200 ng/mL and 10 μL of each sample was then injected.

The extract was fractionated through C18 FlashPure 12g column; flow rate: 30 mL/min into 5 fractions using a gradient of solvents (A) Ammonium bicarbonate buffer pH = 10.2 and (B) ACN (0–5 min 98% A, 5.1–10 min 50% A, 10.1–16 min 40% A, 16.1–27 min 35% A, 27–38 min 35–5% A, 38.1–42 min 5%, 42–46 min 2%. The fractions were collected automatically in 20 mL vials.

The fractions containing yuzurimine were further purified using a semi-preparative column (Waters XBridge C18 5μm column of size 10 x 150 mm) and eluted with a gradient 50-5% ammonium bicarbonate buffer pH = 10.2 and with 6.6 mL/min flow rate acetonitrile controlled manually to yield yuzurimine (15 mg).

Yuzurimine. δ_H_ (400 MHz, CDCl_3_), 5.36 ppm (1H, dd, *J* = 12.0, 7.0 Hz, H-3), 4.40 (1H, d, *J* = 11.5 Hz, H-7a), 4.29 (1H, d, *J* = 11.5 Hz, H-7b), 3.85-3.65 (1H, m, H-12a or H-13a), 3.60 (3H, s, H-25), 3.59-3.47 (1H, m, H-12a or 13a), 3.46-3.32 (1H, m, H-12 or 13b), 3.33-3.16 (1H, m, H-12b), 3.10-2.97 (1H, m, H-20 and H-16a), 2.94-2.80 (1H, m, H-11), 2.79-2.66 (1H, m, H-23), 2.59 (1H, dt, *J* = 7.5, 7.5 Hz, H-1), 2.51 (2H, m, H-15a and H-16a), 2.39-2.32 (2H, m, H-19), 2.06-22.03 (1H, m, H-15b), 2.01 (3H, s, H-9 or H-27), 2.00 (3H, s, H-9 or H-27), 1.92-1.80 (2H, m, H-22), 1.58-1.42 (2H, m, 2), 1.40-1.31 (1H, m, H-18), 1.05 (3H, d, *J* = 7.5 Hz, H-10); δ_C_ (100 MHz, CDCl_3_), 175.4 (C24), 170.9 (C8 or C26), 170.2 (C8 or C26), 72.5 (C3), 66.9 (C7), 57.8 (C12 or C13), 57.6 (C12 or C13), 51.3 (C25), 45.1 (C4), 29.8 (C14), 27.1 (C2), 21.4 (C9 or C27), 21.3 (C9 or C27), 14.8 (C10). NMR data were consistent with those reported in literature [Supplementary Data 5, reference (*28*)].

Similarly, the fractions containing daphniphylline were further purified using a semi-preparative column (Waters XBridge C18 5μm column of size 10 x 150 mm) and eluted with a gradient 50-5% ammonium bicarbonate buffer pH = 10.2 and with 6.6 mL/min flow rate acetonitrile controlled manually to yield daphniphylline (5 mg).

Daphniphylline. δ_H_ (400 MHz, CDCl_3_), 5.63 (1H, dd, *J* = 12.5. 3.0 Hz, H-17), 4.49 (1H, dd, *J* = 13.0, 2.0 Hz, H-24a), 3.72 (1H, d, *J* = 13.0 Hz, H-17b), 3.49-3.29 (1H, m, H-6), 3.24-2.92 (2H, m, H-7), 2.11 (3H, s, H-28), 1.40 (3H, s, H-26), 1.33 (3H, s, H-25 or H-32), 1.01 (3H, d, *J* = 7.0 Hz, H-30 or H-31), 0.99 (3H, d, *J* = 5.0 Hz, H-30 or H-31), 0.91 (3H, s, H-25 or H-32). NMR data were consistent with those reported in literature [Supplementary Data 5, (*29*)].

### Cryosection preparation

Tissue segments (approximately 0.5 cm x 0.5 cm) of petioles, and stems were cut and embedded in 15% gelatin. The tissues were then mounted to a holder using Tissue-Tek O.C.T compound, and sectioned (at −20°C) to a thickness of 40 to 80 μm by Leica CM950 Cryostat (Leica Biosystems). Suitable sections were mounted to SuperFrost Plus adhesion glass slides. The slides were then inspected under bright-field microscopes (Nikon Eclipse E600, and Leica MZ16, Leica microsystems) prior to analysis.

### MALDI-MS analysis

Sections were sprayed with 40 mg mL^-1^ 2,5-dihydroxybenzoic acid matrix in aqueous 50% methanol, 0.1% trifluoroacetic acid by robotic sprayer (TM-sprayer, HTX Technologies, Carrboro, NC). The following parameters were used: flow rate, 50 mL min^-1^; spray nozzle velocity, 1250 mm min^-1^; nozzle temperature, 80°C; track spacing, 3 mm; passes, 24 criss-cross and offset; nitrogen pressure, 10 Psi. Post-spraying, a saturated solution of red phosphorus in acetone was spotted adjacent to the tissue section on each slide.

MALDI-MS acquisition was performed using a Waters SYNAPT G2-Si qTOF mass spectrometer operated in sensitivity or resolution mode with positive ionisation. The instrument was externally calibrated against red phosphorus for each section analysed. MS spectra were acquired between *m/z* 250-600, with the following parameters specified: ScanTime, 0.5s; PlateSpeed, 1.0 s pixel^-1^; Fixed Trap CE, 4.0; Fixed Transfer CE, 2.0; Quad profile, automatic. Spectra were acquired with a 25 μm spatial resolution. Instrument control, data acquisition, and processing were performed using Waters HDI software (v1.5).

The generated MS images represent the spatial intensity distribution of certain *m/z* signals, which can be viewed as an intensity or pseudo-color heat map (with the dark green/black colour being the most intense). For enhanced information recovery and analysis of the different tissue specimens, the data were further processed using python script (BASIS) (*30*). The *m/z* values were binned and aligned to new mass bins common across all the analysed slides. Further intra-sample and inter-sample processing was performed to remove biologically unrelated pixel-to-pixel variation and to adjust for sample-to-sample differences in the overall signal intensity (*30*). The high mass accuracy LC-MS data was used as a reference for the selection of relevant peaks. Alkaloids with the highest log2 value of the abundance across all samples were selected to investigate their distribution using MS imaging.

Linear spatial correlations between the ion images of interest were calculated using HDI software (v1.5), the obtained correlation matrix was used to build cluster heatmap via the pheatmap function in Rstudio using default settings (hierarchical clustering).

### R packages

R analysis was conducted using R-studio (releases 2019-2022). Packages used include tidyverse (*31*), umap (*32*), pheatmap (*33*), fossil (*34*) and FSA (for dunnTest) (*35*).

## Results

### Data preparation

We set out to perform a metabolomics analysis of *Daphniphyllum macropodum* to identify the variation in alkaloids between different individuals and tissue types. Due to limited access to bulk material, we were not able to extract or obtain many chemical standards to use for assigning peaks unambiguously (Supplementary Table 1). Consequently, we developed a workflow that combined MS^1^ *in silico* database annotation, MS^2^ molecular networking and guided clustering to annotate the data holistically (Fig. 2). This method avoids reliance on ambiguous annotations and leverages the large dataset size to identify trends in metabolite distribution.

**Figure 2.**
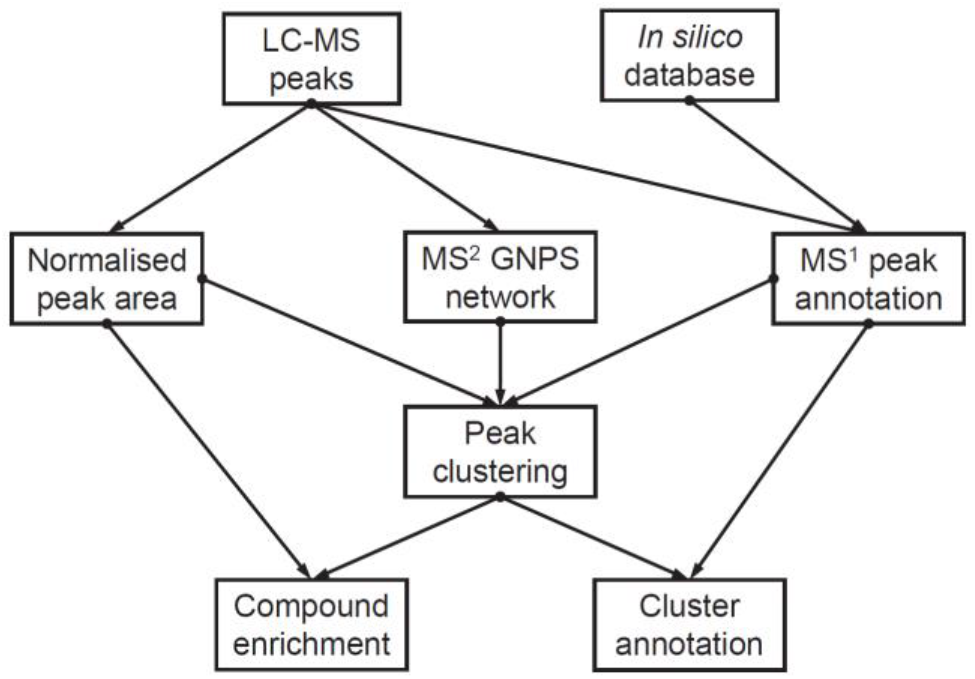
Workflow for analysing untargeted LC-MS data.

We assembled an *in silico* database of *Daphniphyllum* metabolites containing 331 *Daphniphyllum* alkaloids (DAs), split into 25 sub-categories based on the carbon skeleton (*3*). The major observed sub-types are calyciphylline A (61), yuzurimine (60) and yuzurine (48), but the majority of subtypes were represented by five or fewer compounds (Supplementary Fig. 1). We then obtained *D. macropodum* plant material from multiple individuals representing a range of ages, native geographic origin and tissues (Supplementary Table 2, Supplementary Data 1). Untargeted metabolomics analysis of 183 samples provided a total of 1886 filtered peaks, of which 568 were putatively annotated using exact mass comparisons with the *in silico* database. To account for different ionisation responses, peak areas were normalised across all samples.

### Sample Clustering

Due to the complexity of the dataset and the large number of unverified peaks, we used clustering to group similar samples or peaks. To determine the most effective clustering method, we assessed the performance of different clustering parameters compared to groupings based on “ground truth” annotations using the adjusted Rand Index (ARI) (Supplementary Data 2). To handle the zero-inflated, sparse dataset, we examined a multi-step clustering approach, with Principal Component Analysis (PCA) and/or Uniform Manifold Approximation and Projection (UMAP) (*36*) for dimensionality reduction followed by clustering with hierarchical or k-means approaches (*37*). We selected a sample clustering method that favoured clustering samples with the same plant and tissue origin together. The best performing clustering methods based on this metric (i.e. highest ARI compared to plant-tissue annotations) were models combining both PCA and UMAP with either k-means or hierarchical complete clustering (Supplementary Fig. 2, Supplementary Table 4). The sample clustering method with the highest ARI separated the samples into six clusters (Supplementary Fig. 2F).

### Peak clustering

Establishing a “ground truth” for determining the best peak clustering method was complex as few peaks were unambiguously annotated. To obtain annotations that could be used to guide peak clustering, we assumed that structurally similar compounds would cluster together, considered reasonable based on the divergent nature of biosynthetic pathways. We also assumed that molecular networks generated by MS^2^ spectra would be able to group structurally similar compounds together. We generated a molecular network using Global Natural Product Social Molecular Networking (GNPS) (Supplementary Fig. 3) (*24*). Peaks were annotated combining GNPS sub-network assignment and putative compound type peak annotations determined by MS^1^ matches to our *in silico* database (e.g. the annotation DA_10 would be applied to peaks annotated as DAs by MS^1^ within GNPS subnetwork 10). These annotations were used as the “ground truth” to assess clustering methods by ARI (Supplementary Fig. 4A, Supplementary Table 4). The method with the highest ARI for the complete GNPS network annotations employs UMAP and hierarchical complete clustering to provide seven clusters (Supplementary Fig. 4B and C).

### Peak annotation

Furthermore, eight peaks were annotated through comparison to verified standards and twelve peaks tentatively annotated through comparison with literature MS^2^ values (Supplementary Table 5). The GNPS analysis was used to annotate based on library matches and molecular network position, with peaks with a majority of neighbours annotated as DAs reannotated as DAs (Supplementary Fig. 5). A major peak in the dataset, M492.3323T262, is likely to be a novel DA adduct as judged by in source and MS^2^ fragmentation. The C_28_H_45_NO_6_+H peak undergoes neutral loss of 146, corresponding to loss of a rhamnosyl or coumaroyl residue, to leave an alkaloid core of C_22_H_35_NO_2_+H (measured *m/z* = 346.2742, exact *m/z* = 346.2746), a formula that matches four previously reported DAs (Supplementary Fig. 6A and B). In the molecular network, this peak has nine neighbouring peaks each appearing to be different derivatives with the same 346 alkaloid core (Supplementary Fig. 6C). All these peaks were designated DAs of unknown subtype for further analysis. Overall the amendments to the peak annotation increased the number of DA annotations to 322.

### Sample cluster content

Clustering of samples appeared largely to be determined by plant origin, rather than tissue type. For example, sample cluster 5 was exclusive to samples from plant C and D, and sample cluster 4 had a majority plant B samples (Supplementary Fig. 7). Sample cluster 6 contained primarily stem and bark samples, the clearest case of tissue type being a greater influence than plant origins. The contrast between C and D with the rest of the plants may be expected based on their origins in the western Chinese Guizhou province compared to the likely origin of the other plants further east, including Japan. The similarity of E, F and G may be related to their genetic similarity or their relatively young age and similar growing conditions.

### Peak cluster content

Understanding the distribution of annotated peaks across the peak clusters (PCs) is a vital aspect of this untargeted metabolomics approach. First, we assessed the numbers of peaks and their overall type classification in each clusters (Fig. 3A). PC3 contained the most peaks but the majority of these are uncategorised (i.e. unannotated) compounds. PC1 and PC4 had over half their peaks unannotated compounds but also approximately a quarter of the peaks were annotated as DAs, with lower proportions of iridoids and phenylpropanoids. PC2 and PC6 were clearly DA associated, with over three-quarters of their peaks annotated as DAs. PC5 and PC7 contained very low numbers of peaks.

**Figure 3.**
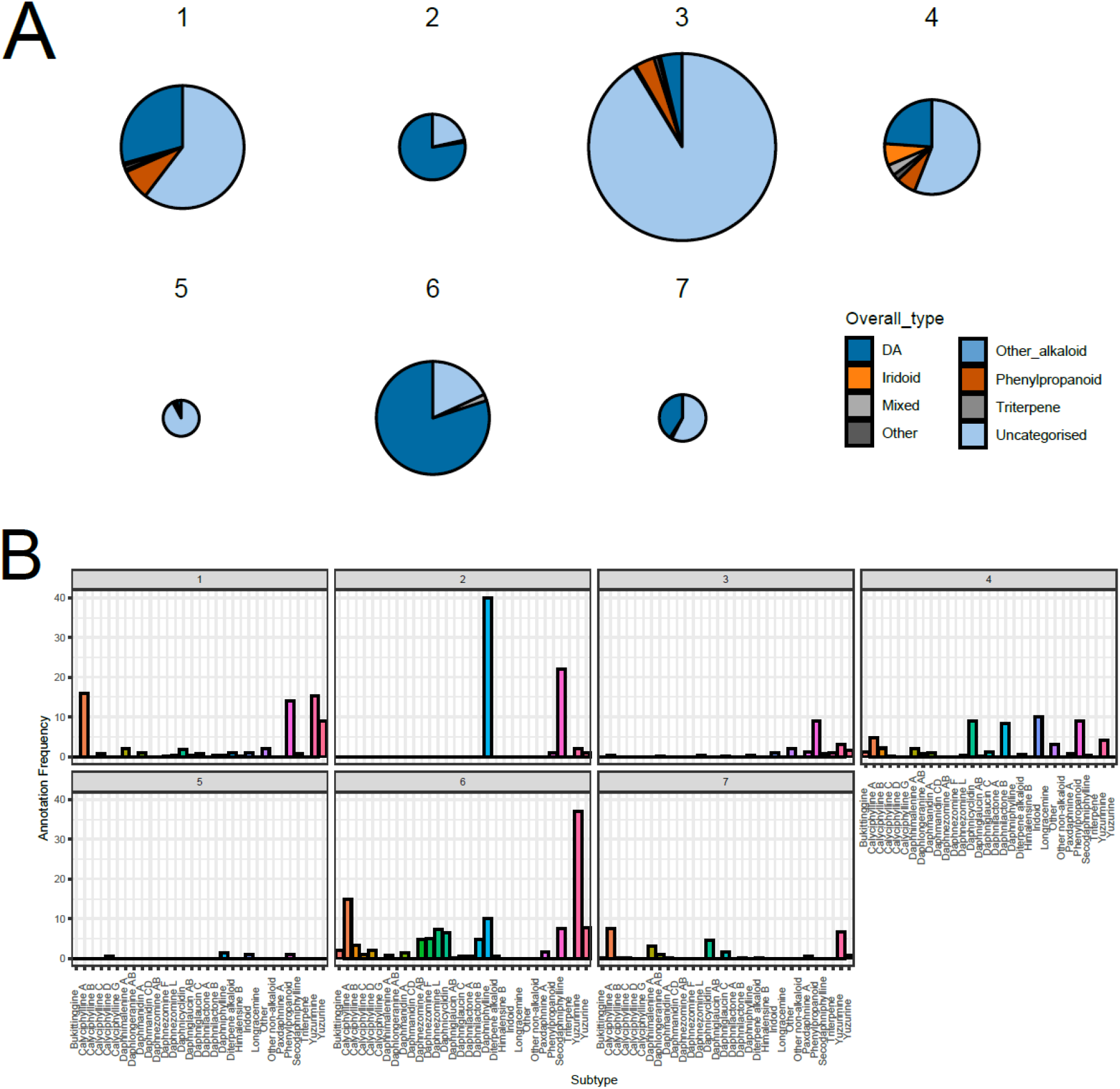
Distribution of annotations in peak clusters. A. Peak type annotations within each peak cluster. The area of each pie is proportional to the total number of peaks in the cluster. B. Peak subtype annotations in each peak cluster, using proportional subtype annotations where multiple annotations are counted as a fraction of a count. Uncategorised peaks not depicted.

We sought to understand the distribution of DA subtypes across the clusters. To do this, we calculated a compound subtype score for each annotated peak based on the proportion of annotations with particular subtypes. These subtype scores were summed within each PC to describe the subtype annotations across the dataset (Fig. 3B).

With this method, a pattern of DA subtype annotation clearly emerges from the peak clusters. Annotated compounds in PC2 are nearly exclusively secodaphniphylline and daphniphylline subtypes. On the other hand, PC6 contains a high proportion of yuzurimine, followed by calyciphylline A, subtype annotations. PC1 has similar distribution to PC6 but with proportionally less yuzurimine subtype. PC4 also contains notable numbers of alkaloid annotations including relatively high daphnicyclidin and daphnilactone B subtypes. The distribution of different subtype annotations in these peak clusters indicates that different alkaloid subtypes accumulate differently in different plants and/or tissues. Determining which plants/tissues accumulate which compounds, coupled to an appreciation of the compound structures, will lead to a greater understanding into their biosynthetic origins.

### Variation in peak clusters across samples

To examine the distribution of PCs across groups we used a metric we termed compound enrichment (CE). We had previously calculated normalised peak values by scaling the data across samples (see above). To gain a single metric to describe a peak cluster or any collection of peaks, we calculated the sum of grouped normalised peak values within each sample. These CE values are then averaged across samples within the sample groups/clusters under examination.

Examining CE in peak clusters across all samples grouped by plant origin, we see significant variation in enrichment across plant origin for all PCs (Fig. 4A, Kruskal-Wallis chi-squared tests, *p* < 0.05, Supplementary Table 6). The DA containing PC1 has increased enrichment in E and F compared to A, B and C (Dunn’s post hoc, *p* < 0.05, Supplementary Table 7 and Supplementary Data 3). PC2 has three statistically significant groupings, with plants either having high (E, F, G), medium (A, B) or low (C, D) enrichment of these compounds. PC6 compounds were found to be either abundant (A, B, E, F, G) or scarce (C, D). CE also varied significantly across sample clusters (Fig. 4B, Kruskal-Wallis chi-squared tests, *p* < 0.05, Supplementary Table 6). Sample cluster 2, which contains an assortment of samples from all plants except C and D, and was significantly enriched in peak cluster 2 compounds compared to all other samples (Dunn’s post hoc, *p* < 0.05, Supplementary Table 8 and Supplementary Data 3).

**Figure 4.**
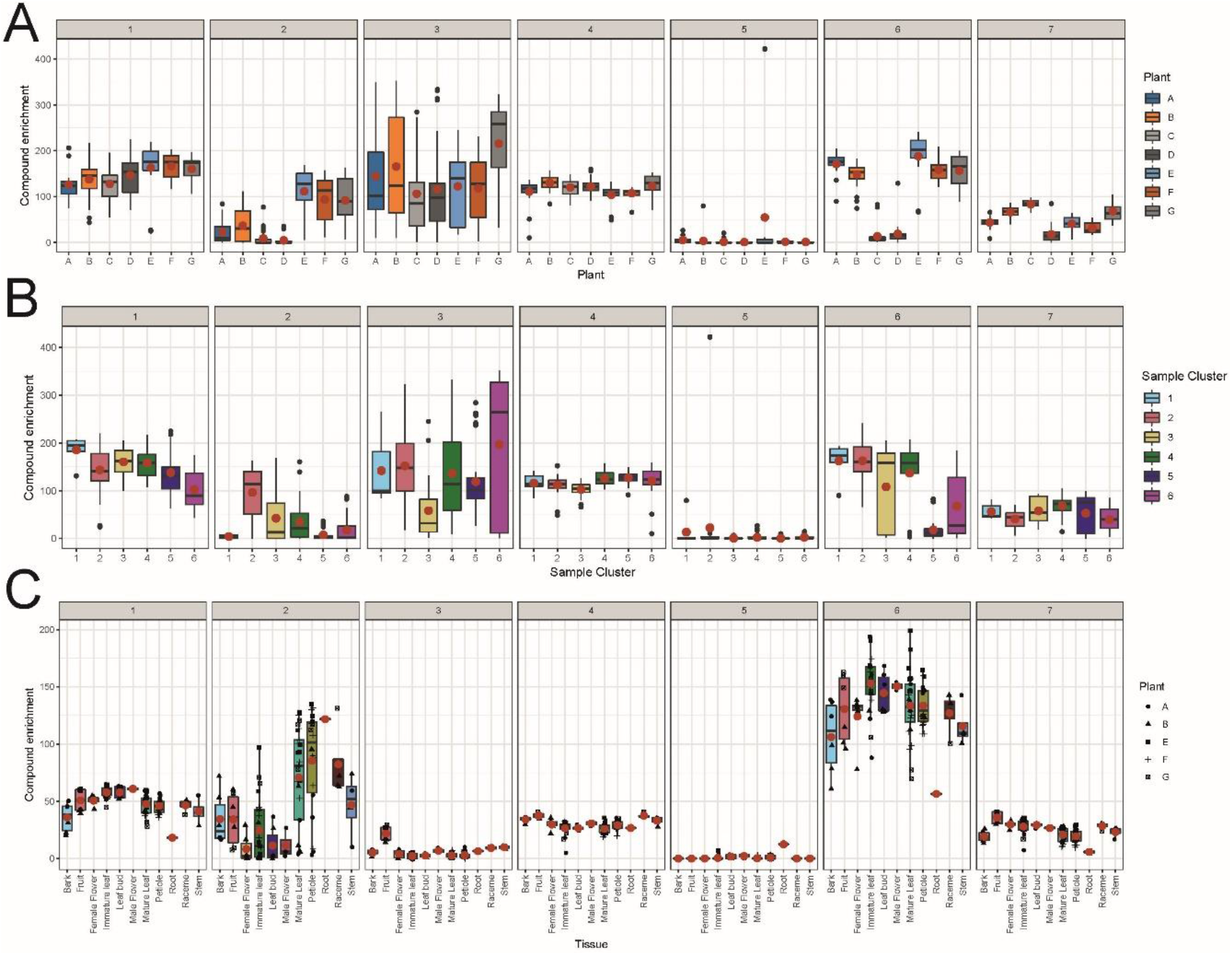
Enrichment of peak clusters across plants (A), sample clusters (B) and tissues (C). Each plot depicts data from peaks associated with specific peak clusters 1 to 7. Compound enrichment is calculated by totalling normalised peak areas within each peak cluster within each sample. Boxplots show distribution of compound enrichment from samples associated with the specific plant (A) or sample cluster (B) or tissue (C). The numbers of samples in each boxplot varies. The means of compound enrichment are shown as a red dot, in (C) plant origin is depicted by point shape and plants C and D were excluded from the analysis. Statistical analysis of differences between groups can be found on Supplementary Tables 6-9 and Supplementary Data 3 and 4.

### Variation in peak clusters across tissues

We next examined the influence of tissue on DA enrichment, initially looking across all plants, though excluding the low DA producers C and D (Fig. 4C). For all peak clusters, CE varied significantly across tissues (Kruskal-Wallis chi-squared tests, *p* < 0.01, Supplementary Table 9). The major DA containing clusters are PC1, PC2 and PC6. Root, bark, stem and petiole samples had relatively low enrichment of PC1, compared to high accumulating samples from immature leaves, leaf buds and male flowers (Dunn’s post hoc test, *p* < 0.05, Supplementary Data 4). The signal from peak cluster 6 was similar, with bark, stem, root, petiole and mature leaf having significantly lower enrichment than immature leaf. PC2 shows a very different pattern: samples with high enrichment are mature leaf, petiole, raceme and root which contrast with the low accumulating leaf buds, immature leaves and female flowers.

PCs 3, 4, 5 and 7 also contain DAs. PC3 and PC5 are enriched in fruit and root respectively. PC4 is enriched in bark, fruit, raceme and stem relative to leaves and root. PC7 is different, with low enrichment in bark, root and petiole, and high in fruit and leaves. We also looked at tissues within individual plants and across different PCs (Supplementary Fig. 8); this largely confirmed the analysis of the bulked data, with all PC enrichment influenced by tissue type in a subset of plants (Kruskal-Wallis chi-squared tests, *p* < 0.01, Supplementary Data 4).

### Summary of peak cluster distribution

Based on CE across plants and tissues, and integrating inferred peak cluster content from annotations, a pattern of DA distributions begins to emerge (Fig. 5). Broadly, plants C and D are relatively low in DAs, but the DAs that do accumulate appear to be from PC1, largely calciphylline A, yuzurimine and yuzurine subtypes, are most abundant in non-vascular aerial tissues. Alkaloids from PC4, daphnicyclidin and daphnilactone B subtypes, accumulate in vascular tissue from C and D (stem and bark). Plants A, B, E, F and G all accumulate yuzurimine subtype PC6 DAs across all tissues, but greatest enrichment is in young non-vascular tissues (i.e. immature leaves, leaf buds). In contrast, PC2 compounds, associated strongly with daphniphylline and secodaphniphylline subtypes, are observed in plant B, E, F and G but primarily in vascular containing tissue and mature leaves.

**Figure 5.**
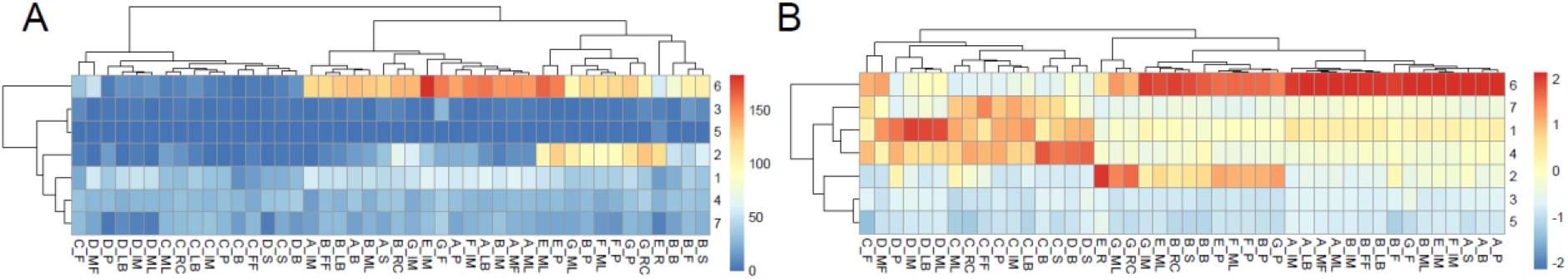
Pattern of DA distribution across plants and tissues. Rows represent peak clusters, columns are plant-tissue annotations. In (A) values are averages of peak cluster compound enrichment in each annotation. In (B) the columns have been scaled and centred so values are z-scores of mean compound enrichment within each plant-tissue annotation. Panel A highlights in what annotation are peak clusters relatively enriched, giving an indication of which tissue accumulate most DAs. Through normalisation, panel B describes DA subtype distribution within each plant-tissue annotation. Only peaks annotated as DAs are included in the analysis. Rows and columns have been sorted by hierarchical clustering. Annotations show plant origin (A-G) followed by tissue type: IM (immature leaf), ML (mature leaf), P (petiole), S (stem), R (root), B (bark), RC (raceme), LB (leaf bud), MF (male flower) and FF (female flower).

Overall, the plant E root samples appear to demonstrate the most atypical pattern being the only samples with higher enrichment for daphniphylline and secodaphniphylline subtypes (peak cluster 2) compared to any other DAs. Furthermore, it also contains relatively high levels of M544.3635T209 from PC 5, predicted to be a daphniphylline subtype compound. This provides evidence that roots contain a different biosynthetic repertoire than aerial tissues. However it must be noted that in this analysis we were unfortunately limited to two root samples from a single plant, and further experiments on root are required to understand DA distribution in roots further.

The results here provides evidence that two factors that influence DA composition: plant origin and organ. Across all tissues, plant C and D produce a distinct set of alkaloids compared to the other plants (A, B, E, F and G). Within each plant, there is a distinct difference between DA subtypes in older and more vascular tissue, compared to younger and non-vascular tissue.

### Imaging mass spectrometry

To investigate the spatial distribution of DAs within specific tissues, we used matrix-assisted laser desorption/ionisation MS (MALDI-MS). For this analysis, cryosections of tissues from plant F were embedded in a matrix. We tested α-cyano-4-hydroxycinnamic acid and 2,5-dihydroxybenzoic acid matrices and determined the latter provided a better response for our target molecules, with DAs detected as protonated ions in positive ion mode using a spatial resolution of 25 μm.

We focused on representative alkaloids that were highly abundant in plant F according to the LC-MS analysis. For the annotation of the selected alkaloids, the mass to charge value accuracy was calculated as mass accuracy (ppm) of the binned and aligned measurements across multiple samples, and a delta ppm of ±17 was used to match to putative molecular formula (Supplementary Table 10). Where there are multiple DAs with the same molecular formula in our *in silico* DA database, we assumed that the MALDI detected DAs corresponded to the most abundant DAs with the same formula in the plant F LC-MS data (Supplementary Fig. 9).

The MALDI-MS analysis of a petiole cross section revealed distinct spatial distribution of DAs, with alkaloids detected with high intensity in the epidermis, phloem region, and the medulla (Fig. 6A-D). A subset of ions, including *m/z* 370.2333 and *m/z* 372.2513 were detected in the epidermis but largely absent in the phloem and medulla (Fig. 6B). In contrast, other ions such *m/z* 470.3621 and *m/z* 486.3571, were readily detected with high intensity in phloem, medulla and epidermis (Fig. 6C). A third pattern was also observed for alkaloids including *m/z* 512.3707 and 528.3670, these were most abundant in the phloem region and medulla whilst largely absent in the epidermis (Fig. 6D). Through matching MALDI-MS ions with LC-MS peaks, it appeared that compounds belonging to PC1 and PC6 seem to localise primarily in the epidermis (Table 1). However, compounds belonging to PC2 appear to localise around the phloem region and medulla and more rarely the epidermis.

**Figure 6.**
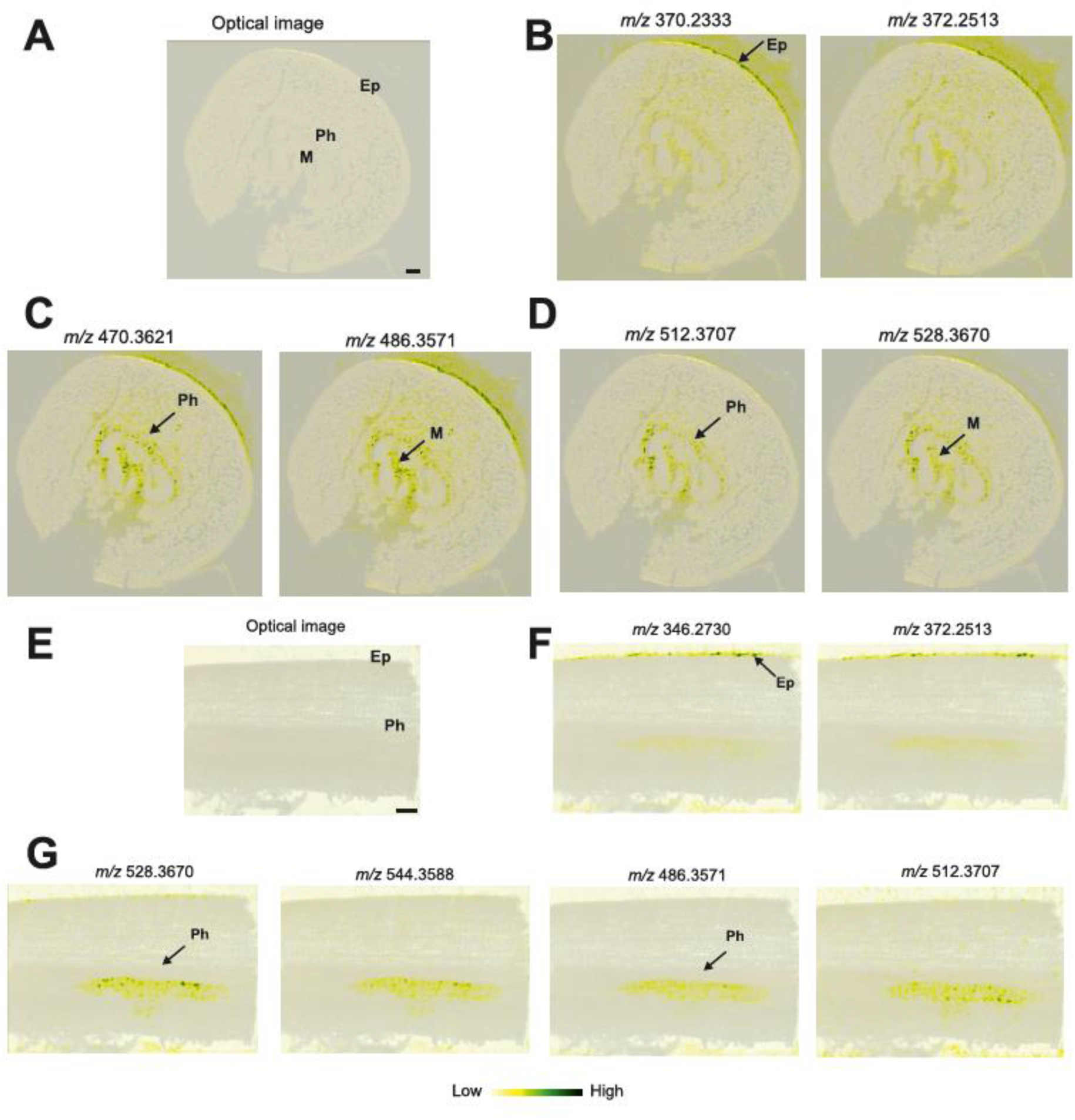
Distribution of DAs within petiole and stem tissue. Ion intensity maps of selected alkaloids detected as [M+H]^+^ ions with the MALDI-MS in the cross section of *D. macropodum* petiole (A-D) and longitudinal section of *D. macropodum* stem (E-G). Micrographs included for comparison (A and E, size bar = 200 μm), which are also added as background to the MS-images. Colour bar represents MS signal intensity. Positions of the epidermis (Ep), the phloem region (Ph), and the medulla (M) are indicated. The different panels indicate patterns of DA distributions with enrichment in the epidermis (B and F), phloem/medulla (D and G) or all locations (C). MALDI-MS analysis of a stem cross section provided similar results (Supplementary Fig. 10). See Table 1 for collected data. Petiole cross section analysed in sensitivity mode (laser energy 80.0 J cm^-2^). Stem longitudinal section analysed in resolution mode (laser energy 250.0 J cm^-2^).

**Table 1.** Localisation of DAs in plant tissue. Presence of DA in region determined by ion intensity map (see Fig. 6 and Supplementary Fig. 10). Compounds were abundant (✓✓), present (✓) or absent (blank). LC-MS peak IDs were determined by matching mass to the compound of highest abundance in plant F (see Supplementary Fig. 9). PC = peak cluster, Ep = epidermis, Ph = phloem.

| MALDI | LC-MS | PC | Petiole |  | Stem 1 |  | Stem 2 |  |
| --- | --- | --- | --- | --- | --- | --- | --- | --- |
| Measured *m/z* | Peak ID |  | Ep | Ph | Ep | Ph | Ep | Ph |
| 342.2426 | M342.2429T159 | 6 | ✓✓ | ✓ | ✓✓ | ✓ | ✓✓ |  |
| 346.2730 | M346.2742T164 | 6 |  |  | ✓✓ | ✓ | ✓✓ |  |
| 360.2573 | M360.2899T261 | 6 |  |  |  | ✓ | ✓✓ |  |
| 362.2674 | M362.2691T83 | 6 |  |  |  | ✓ | ✓✓ |  |
| 368.217 | M368.2222T191 | 1 | ✓✓ | ✓ | ✓✓ |  | ✓✓ |  |
| 370.2333 | M370.2378T179 | 6 | ✓✓ | ✓ | ✓✓ |  | ✓✓ |  |
| 372.2513 | M372.2539T131 | 6 | ✓✓ | ✓ | ✓✓ | ✓ | ✓✓ |  |
| 374.2678 | M374.2691T120 | 6 | ✓✓ | ✓ | ✓ | ✓ | ✓✓ |  |
| 376.2856 | M376.2848T264 | 6 | ✓✓ | ✓ |  |  | ✓✓ | ✓✓ |
| 386.2364 | M386.2328T206 | 1 | ✓✓ |  |  |  | ✓✓ |  |
| 414.2611 | M414.2641T208 | 2 | ✓✓ | ✓ | ✓ | ✓ | ✓✓ | ✓ |
| 470.3621 | M470.3631T259 | 2 | ✓ | ✓ | ✓ | ✓✓ |  | ✓✓ |
| 472.2658 | M472.2695T209 | 6 | ✓✓ | ✓ | ✓ | ✓ | ✓✓ | ✓ |
| 472.3722 | M472.3787T248 | 2 | ✓ | ✓✓ |  | ✓✓ |  | ✓✓ |
| 484.3440 | M484.3424T213 | 2 |  | ✓✓ |  |  | ✓✓ | ✓✓ |
| 486.3571 | M486.3581T252 | 2 | ✓✓ | ✓✓ |  | ✓✓ | ✓✓ | ✓✓ |
| 488.3658 | M488.3738T267 | 2 | ✓ | ✓ |  | ✓✓ | ✓✓ | ✓✓ |
| 502.3462 | M502.3529T221 | 2 | ✓ | ✓ |  | ✓✓ | ✓ | ✓ |
| 512.3707 | M512.3737T250 | 2 |  | ✓✓ |  | ✓✓ |  | ✓✓ |
| 528.3670 | M528.3688T268 | 2 |  | ✓✓ |  | ✓✓ |  | ✓✓ |
| 544.3588 | M544.3635T260 | 2 |  | ✓✓ |  | ✓✓ |  | ✓✓ |

Further examination of alkaloid localisation in a longitudinal stem section showed similar trends, with the alkaloids detected in the epidermal cells and the phloem region (Fig. 6E-G). For instance, the molecular ions *m/z* 346.2730 and *m/z* 372.2513 were distributed mostly in the epidermal cells and to a lesser extent in the phloem. Molecular ions including *m/z* 472.2658, 486.3571, 528.3670 and 544.3588 were detected mostly in the phloem region. This result validates again the same observation, that alkaloids of PC1 and PC6 are localised in the epidermal cells, while alkaloids belonging to PC2 tend to localise primarily in the phloem region. Further stem cross sections showed similar distribution patterns (Supplementary Fig. 10).

For each MS image, we generated spatial correlation values and a heatmap, which highlighted groups of spatially correlated compounds (Fig. 7). When we inspected the annotation of these alkaloids, we noticed that compounds that belong to PC1 and PC6, indicating co-localisation in the analysed cross-sections (e.g. *m/z* 342.2426, 368.217 and 372.2513). Alkaloids that belong to PC2, such as *m/z* 544.3588, 512.3707, and 528.3670, seemed to spatially co-localise. This further confirmed our visual observation that alkaloids in the same peak clusters tend to have similar spatial distribution within tissues.

**Figure 7.**
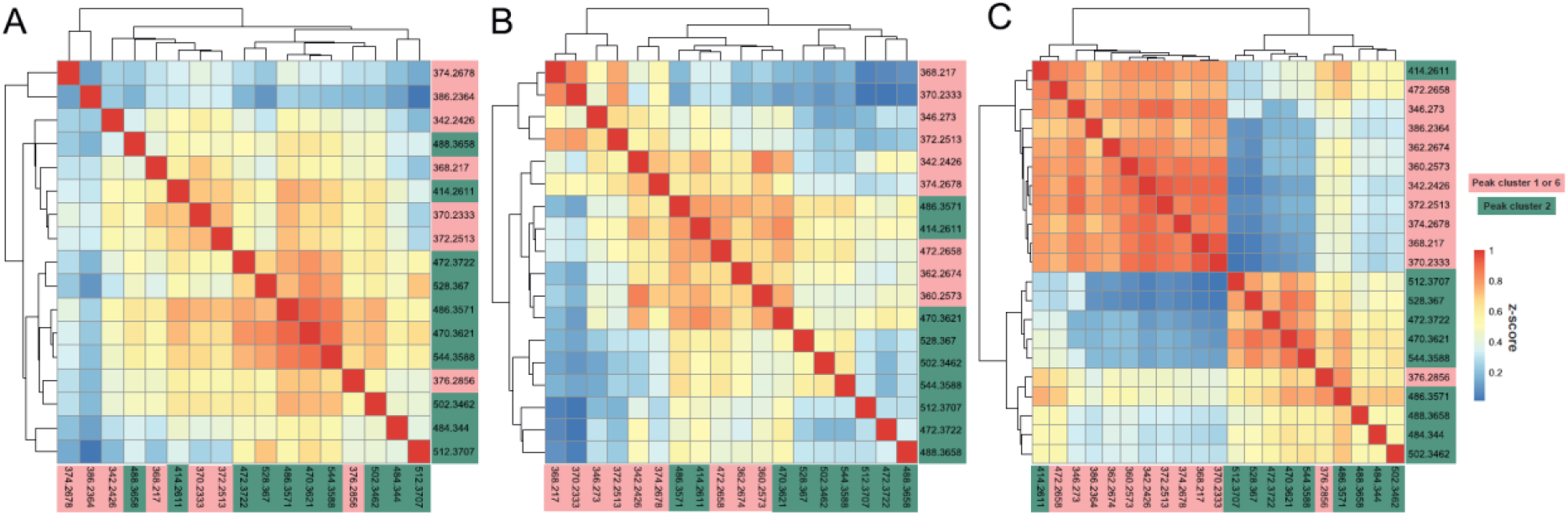
Spatial correlations of selected DAs within *D. macropodum* tissues. A. Petiole cross section (images on Fig 6A-D). B. Longitudinal cross section (images Fig. 6E-G). C. Stem cross section (images Supplementary Fig. 10). See Table 1 for further details.

The spatial distribution of DA subtypes within tissues determined by MALDI-MS mirrors patterns of distribution between tissues determined by LC-MS. In LC-MS analysis we found PC2 compounds, of daphniphylline and secodaphniphylline subtype, associated with organs containing vascular tissue, whilst yuzurimine subtype PC6 compounds associated with younger, low-vascular tissue containing organs. Accordingly, the MALDI-MS images shows ions corresponding to PC2 DAs associated with the vascular phloem tissue, whereas ions representing PC6 DAs are enriched in the epidermis. A small subset of ions corresponding to PC2 DAs were found in both tissues, likely a result of dual location. Overall, the agreement between the independently conducted MALDI-MS and LC-MS experiments verifies that DA subtypes demonstrate distinct distribution in plants.

## Discussion

The alkaloids from *Daphniphyllum* interest researchers as examples of natural products with structural complexity and skeletal diversity. They have also inspired total synthesis campaigns where cutting-edge chemical methods alongside ingenious synthetic steps attempt to replicate nature’s biosynthetic feats (*9*). There has been some, if limited, examination of DA bioactivities (*2*). Aside from early labelling experiments (*16*) and synthetic biomimetic syntheses (*10*), a biological or biosynthetic examination of the DA alkaloids is lacking. Our ultimate aim is to understand how *Daphniphyllum* biosynthesises DAs. In the process of understanding the pathway, we hope to obtain enzymes that will allow us gain access to DA in heterologous systems, allowing facile isolation followed by bioactivity testing (*38*).

### LC-MS Workflow

The first step of this long-term project was to gain an understanding of the distribution of DAs across different available plants, their organs and tissues using metabolomics approaches, namely LC-MS and MALDI-MS. A major challenge of working in this new system was the lack of available standards. This was largely a result of us lacking sufficient plant material from which to isolate compounds. Commercially purchased DA standards did not pass our quality control checks. *In lieu* of chemical standards, we developed a new workflow which combined MS^1^ annotations, MS^2^ networking and clustering (Fig. 2). This method enabled us analyse the large LC-MS dataset holistically without relying on ambiguous annotations of single peaks but instead using cumulative annotations across peak clusters. This workflow and clustering method can be generalised to other metabolomics datasets and even other experimental types where the data have a similar structure.

### LC-MS summary

The LC-MS analysis revealed DA subtypes vary across plants and organs. Plants C and D, originally from Guizhou province, China, produce daphnicyclidin and daphnilactone B subtypes in stem and bark, and calciphylline A, yuzurimine and yuzurine subtypes in leaves and flowers. The other plants investigated here, either originating from Japan or horticultural sources (original geographic origin unknown, possibly Japan), accumulate yuzurimine subtypes primarily in younger leaves, whilst (seco)daphniphylline subtype compounds accumulate more in organs containing vascular tissue. Clearly genetic origin plays a role in DA content and distribution as plants C and D are very closely related (probable siblings) compared to the other plants, yet are grow in the UK in the same location as plants A and B, ruling out purely environmental factors. It is notable that the plants described as the Japanese low growing *humile* variant of *D. macropodum* (B and G) did not differentiate themselves metabolically from the horticultural derived *D. macropodum* (A, E and F), perhaps confirming *humile*’s status as a variant of *D. macropodum* rather than a sub- or separate species (*39*).

The results here match with studies into *D. oldhamii*. Two *D. oldhamii* plants from different regions of Guizhou province were found to produce different subtypes DAs, with yuzurimine and daphnicyclidin subtypes showing differences between the individuals (*40*). Collation of data from natural product isolations of *D. oldhamii* also indicates DA subtypes are differentially distributed in organs, with yuzurimine subtypes primarily in leaves, (seco)daphniphylline subtypes in root and yuzurine subtypes in fruit (*20*).

### MALDI-MS

MALDI-MS imaging can complement metabolomic analysis of homogenised tissues in LC-MS (*41*). The ion intensity maps showed that DAs were localised in specific regions of the petiole and stem tissues. Correspondence of MALDI-MS ions with LC-MS peaks provided access to peak cluster/subtype information. The independently acquired MALDI-MS data validated the LC-MS work, with (seco)- daphniphylline subtypes associated with vascular tissue (phloem) and yuzurimine subtypes localised to the epidermis.

### Epidermal localisation

In ecological context alkaloid localisation within epidermal cells may suggest a role in defence, as the epidermis is the most exposed tissue to environmental stressors such as pathogens and herbivores. Epidermal cells have been found to accumulate alkaloids in other plants, such as quinolizidine alkaloids in lupin species (*42*), where they act to deter herbivores (*43*). There is evidence that some DAs have insecticidal (*44*) or pesticidal (*45, 46*) bioactivities, though this has not been thoroughly explored. The yuzurimine subtype compounds that accumulate in the epidermis are also significantly enriched in immature leaf, a tissue that is particularly at risk of attack as they lack the waxy protective layer of mature leaves. Altogether, the localisation of yuzurimine subtype DAs in the epidermis, and accumulation in immature leaf, strongly suggests their role in defence against bioagressors.

### Phloem localisation

DAs from secodaphniphylline and daphniphylline subtypes appear to be enriched in organs containing mature vascular tissue, and, correspondingly, are found to localise to tissue in the phloem region. We propose two hypotheses for phloem-localisation: transportation or specialisation. First, it is possible that DAs are being transported through the phloem to other parts of the plant: it has been shown that alkaloids can be transported through vascular tissues from source to sink organs. For instance, in some *Nicotiana* species, nicotine is biosynthesised in the root tissues and is translocated to the leaves via xylem transport (*47*). Similarly, it has been shown that pyrrolizidine alkaloids from *Senecio vulgaris* are translocated via the phloem from the roots, where they are synthesised, into sink organs such as flowers or seeds (*48, 49*).

The alternative is that there may be specialised cells near the phloem region where alkaloids are being produced and/or stored. Plants often accumulate natural products in specific specialised cells, to limit interference with active biosynthesis and possible cytotoxic effects (*38*). In opium poppy for instance, benzylisoquinoline alkaloids accumulate in the latex of specialised cells associated with the phloem throughout the plant, known as laticifers (*50*). Internal secretory cells are often associated with vascular tissues, and commonly observed in tropical habitat (*51*). However, the examination of our *D. macropodum* specimens to identify internal secretory elements around the phloem region proved to be inconclusive and additional literature search showed that their presence was debated (*52, 53*).

### Transport

The involvement of multiple organs and tissues in a complex alkaloid pathway is reminiscent of both benzylisoquinoline alkaloid (BIA) and monoterpene indole alkaloid (MIA) biosynthesis. Immunolabeling and *in situ* hybridisation in alkaloid producing plants such as *Catharanthus roseus* (MIAs) and opium poppy (BIAs) show that alkaloid accumulation site is not necessarily the synthesis site, and that often we see the involvement of diverse cell types in the biosynthesis (*54*). For instance, *C. roseus* divides alkaloid biosynthesis among internal phloem-associated parenchyma, epidermal cells, laticifers, and idioblasts (*55*). MS imaging investigations of *C. roseus* revealed that MIAs and their precursors in the plant have different spatial distributions. These studies demonstrated that most MIA iridoid precursors are localised in the epidermal cells, while major MIAs including serpentine and vindoline are localised in idioblast cells which suggest the involvement of transport processes (*56, 57*). Similar phenomena may be occurring in *Daphniphyllum*, with biosynthetic pathways split across different tissues with compound transport connecting them.

### Biosynthetic pathway

By considering the biosynthetic pathway alongside metabolite localisation we can build a speculative model of DA biosynthesis in the plant. It must be noted that the number of carbons in the core skeleton of DAs is either C30 or C22, with C30 compounds are exclusively found within the daphniphylline or secodaphniphylline subtypes. As all DAs are derived from squalene (C30), and likely pass through a single entry-point, all C22 DAs are expected to derive from one or more C30 DAs. Evidence of the pathway origin is found in the seminal syntheses by Heathcock (*10, 15*), which place the secodaphniphylline as the core skeletal intermediate that can lead to all other subtypes. Notably, daphniphylline subtypes are not thought to be intermediates but end products in their own right (*20*).

Either the epidermally localised DAs are synthesised *de novo in situ* or there is transportation of secodaphniphylline-like precursors from phloem region (Fig. 8A). Interestingly the MALDI-MS ion *m/z* 470.3621 (LC-MS: M470.3631T259, PC2) matches the mass of secodaphniphylline and is found in both phloem and epidermal regions (Table 1). The ion *m/z* 346.2730 (LC-MS: M346.2742T164, PC6) matches daphnezomine M, a C22 secodaphniphylline previously reported in *D. macropodum*, and is enriched in the epidermis (Table 1) (*58*). These two compounds, or closely related compounds, may be the link between the phloem and epidermis and the transition of C30 to C22 DAs (Fig. 8A). A similar question arises for phloem localised DAs: are these product of transportation or are they formed *in situ* (Fig. 8B)? Whilst we have limited data for root, it is the organ most enriched in (seco)daphniphylline subtypes and may be the source of early-pathway compounds. Major root peaks include M528.3688T230 and M544.3635T209, which may be daphnezomine D and daphnezomine E respectively based on previous reports in *D. macropodum* (*59*). Curiously, these compounds are N-oxides, a chemical form that has been proposed to be crucial for allocation of pyrrolizidine alkaloids from source to sink via phloem, as they cannot pass through membranes (*60, 61*).

**Figure 8.**
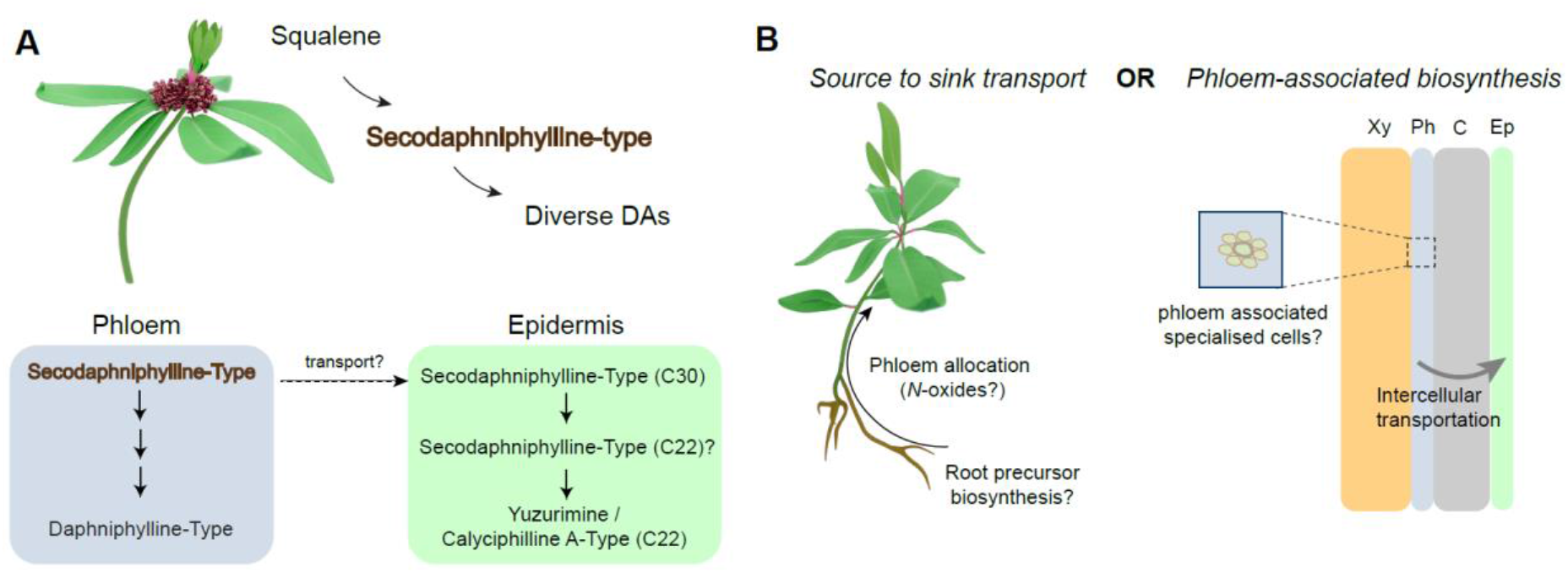
Hypotheses for DA biosynthesis across different tissues. A. Tissue distribution of DAs with hypothesised transportation of secodaphniphylline precursors from phloem to epidermis. B. Origin of phloem located secodaphniphylline precursors, either from root or specialised cells. Xy = xylem, Ph = phloem, C = cortex, Ep = epidermis.

## Conclusion

We have employed MALDI-MS and LC-MS to provide clues about the biological function and biosynthetic origins of *Daphniphyllum* alkaloids. Our analysis indicates that alkaloid distribution within the tissues is complex, with the possible involvement of intercellular and source to sink transport, or specialised cells. *Daphniphyllum* alkaloids have a remarkable skeletal diversity rarely seen in other plants, and the structure dependent distribution discovered here represents a significant step forward in understanding their organisation and origin. The gained insights on the distribution of alkaloids in *Daphniphyllum* can guide further gene discovery efforts through examining gene expression across different cell and tissue types through, for instance, single cell RNAseq (*62*). Through combining metabolomics and genomic approaches we aim to understand how *Daphniphyllum* constructs molecules of remarkable complexity and variety.

## Acknowledgements

We acknowledge the invaluable contributions of the University of York Biology Technology Facility, the Centre of Excellence in Mass Spectrometry, the Biology Horticultural Facility and the Chemistry NMR facility in conducting this research. We thank Karen Hogg and Clare Steele-King in the Biology Technology Facility for support with mass-spectrometry imaging sample preparation. We thank Kotaro Yamamoto for early stage discussions on mass-spectrometry imaging and Olivier Leroux for advice on microscopy. We thank the Yorkshire Arboretum and Castle Howard for access to the plants on their site. In particular we thank John Grimshaw for overseeing the collaboration and Jonathan Burton for arranging site access. We are very grateful for the support and horticultural advice from Sue and Bleddyn Wynn-Jones, including aid with taxonomic assignment. B.R.L., K.E. and the overall *Daphniphyllum* project are supported through a UKRI Future Leaders Fellowship awarded to B.R.L (MR/S01862X/1). B.R. is supported by a BBSRC White Rose DTP studentship (BB/T007222/1). We acknowledge a University of York Research Priming award for funding S.C.

