## Supplementary material for "Integrative metabolomics reveal the organisation of alkaloid biosynthesis in *Daphniphyllum macropodum*": 2_iterateClusters_annotated.R

```
setwd("C:/WD/") #### Direct this to your working directory
library(fossil)
library(umap)
library(Spectrum)
set.seed(1)
dataFile="dataFile.csv"    ####This is your dataFile
annotFile="AnnoFile.csv"   ####This is your annotation file (with ground truth)
outputFile="output.csv"  ####This is your output file
a=read.csv(dataFile, stringsAsFactors = F, header=T, row.names = 1)
data=t(apply(a, c(0,1), function(i){as.numeric(i)}))                     #####remove t() to make it work on non-transposed data
annot=read.csv(annotFile, stringsAsFactors = T, header=T, row.names = 1)

umapFun <- function(data, pcs=0){
    #no UMAP
    clusterFun(data, pcs)
    #UMAP
    sapply(seq(2,min(16,(dim(data)[2]-1)),2), function(comp){       ####range of UMAP components
        sapply(seq(5,20, 2), function(neighbours){                  ####range of UMAP neighbours
            custom.settings = umap.defaults
            custom.settings$n_neighbors = neighbours
            custom.settings$n_components = comp
            out=umap(data, config=custom.settings)$layout
            clusterFun(out, pcs, comp, neighbours)
        })
    })
}

calculateStats<- function(cluster, annot, pcs, comp=0, neighbors=0, algorithm){
    
    if(max(table(cluster))/length(cluster)<0.5){
        
        
        print(length(as.numeric(as.factor(cluster))))
        print(length(as.numeric(as.factor(annot[,1]))))
        
        #calculate Rand Index
        rand=rand.index(as.numeric(as.factor(cluster)), as.numeric(as.factor(annot[,1])))
        print(rand)
        
        #calculate adjusted Rand Index
        adjRand=adj.rand.index(as.numeric(as.factor(cluster)), as.numeric(as.factor(annot[,1])))
        print(adjRand)
        #calculate True positive rate for each type
        categories=table(annot[,1])
        categories=categories[which(categories>1)]
        ncat=names(categories)
        tpr=sapply(ncat, function(i){
            sub=cluster[which(as.character(annot[,1])==i)]
            (sum(table(sub)*table(sub))-length(sub))/(length(sub)*length(sub)-length(sub))
        })
        names(tpr)=ncat
        
        
        
        #calculate the same thing, but removing any uncategorised values:
        
        print(paste(annot[,1])[1:3])
        
        select.all.minus.uncat <- which(paste(annot[,1]) != "Uncategorised") # readded.
        cluster.red <- cluster[select.all.minus.uncat]
        annot.red <- annot[select.all.minus.uncat,1]
        rand.red <- rand.index(as.numeric(as.factor(cluster.red)), as.numeric(as.factor(annot.red)))
        #print(rand.red)
        adjRand.red <- adj.rand.index(as.numeric(as.factor(cluster.red)), as.numeric(as.factor(annot.red)))
        #print(adjRand.red) # TPR too removing uncategorised:
        categories.red <- table(annot.red)
        categories.red <- categories.red[which(categories.red>1)] # OK here?
        ncat.red <- names(categories.red)
        #print(as.character(annot.red))
        #tpr.red <- sapply(ncat.red, function(i){
        #    sub.red=cluster.red[which(as.character(annot.red)==i)]
        #   (sum(table(sub.red)*table(sub.red))-length(sub.red))/(length(sub.red)*length(sub.red)-length(sub.red))
        #})
        #names(tpr.red) <- ncat.red
        
        
        if(!file.exists(outputFile)){
            colNames=paste(c("usePCA?", "numPCs", "useUMAP?", "numNeighbours", "numadjsComponents", "clusteringAlgorithm", "K", "RandIndex", "adjustedRandIndex", "RandIndex:no_uncategorised", "adjustedRandIndex:no_uncategorised",ncat, rownames(annot)), sep=",", collapse =",")
            
            write(colNames, outputFile)
        }
        
        line=paste(c(pcs>0, pcs, comp>0, neighbors, comp, algorithm, length(unique(cluster)), rand, adjRand, rand.red, adjRand.red, tpr, cluster), sep=",", collapse =",")
        write(line,file=outputFile,append=TRUE)
    }
}    

clusterFun <- function(data, pcs=0, comp=0, neighbors=0, consensusOnly=F){
    if(!consensusOnly){
        #k-means
        sapply(c(5:15), function(K){
            clusters=kmeans(data, K, nstart=20)$cluster
            calculateStats(clusters, annot, pcs, comp, neighbors, "K-means")
        })
        
        #hierarchical clustering
        sapply(c("complete", "median"), function(algo){  #removed "average" for speed
            temp=hclust(dist(data), method=algo)
            sapply(c(5:15), function(K){
                clusters=cutree(temp, K)
                calculateStats(clusters, annot, pcs, comp, neighbors, paste("hierarchical", algo, sep="_"))
            })
        })
    }
 ####Uncomment the below to add spectral clustering
 #   #spectral clustering
 #   if(pcs==0 || comp>0){
 #       if(!consensusOnly){
 #           a=Spectrum(as.data.frame(t(data)), maxk = 15, method = 2, showpca=F, tunekernel=TRUE)
 #           clusters=a[[1]]
 #           calculateStats(clusters, annot, pcs, comp, neighbors, "spectral")
 #       }else{
 #           mats=sapply(c(1:5), function(cons){
 #               a=Spectrum(as.data.frame(t(data)), maxk = 15, method = 2, showpca=F, tunekernel=TRUE)
 #               a[[1]]
 #           })
 #           matsTogether=Reduce('+', mats)
 #           sapply(2:15, function(K){
 #               clusters=kmeans(matsTogether, K=K, nstart=20)$cluster
 #               calculateStats(clusters, annot, pcs, comp, neighbors, "consensus_spectral")
 #           })
 #           
 #       }
 #   }
    
}

####################You might want to run this first (testing without PCA)
# no PCA
  umapFun(data)
  
  
##################And then you can run this (testing with PCA)
outputFile="OutputwithPCA.csv"  ####This is complementary output
  
#PCA
sapply(3:10, function(pcs){  #range of number of PCs to include
    dataPC=prcomp(data)$x[,1:pcs]
    umapFun(dataPC, pcs)
})    
    
    
    
    
```
