## Supplementary material for "Integrative metabolomics reveal the organisation of alkaloid biosynthesis in *Daphniphyllum macropodum*": 5_NMR_data.pdf

### Yuzurimine

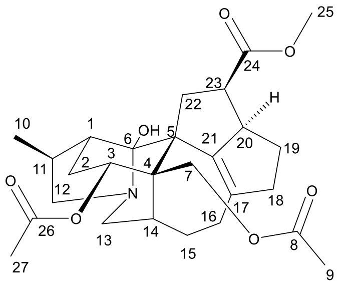

$\delta_H$ (400 MHz, $CDCl_3$), 5.36 ppm (1H, dd, $J$ = 12.0, 7.0 Hz, H-3), 4.40 (1H, d, $J$ = 11.5 Hz, H-7a), 4.29 (1H, d, $J$ = 11.5 Hz, H-7b), 3.85-3.65 (1H, m, H-12a or H-13a), 3.60 (3H, s, H-25), 3.59-3.47 (1H, m, H-12a or 13a), 3.46-3.32 (1H, m, H-12 or 13b), 3.33-3.16 (1H, m, H-12b), 3.10-2.97 (1H, m, H-20 and H-16a), 2.94-2.80 (1H, m, H-11), 2.79-2.66 (1H, m, H-23), 2.59 (1H, dt, $J$ = 7.5, 7.5 Hz, H-1), 2.51 (2H, m, H-15a and H-16a), 2.39-2.32 (2H, m, H-19), 2.06-22.03 (1H, m, H-15b), 2.01 (3H, s, H-9 or H-27), 2.00 (3H, s, H-9 or H-27), 1.92-1.80 (2H, m, H-22), 1.58-1.42 (2H, m, 2), 1.40-1.31 (1H, m, H-18), 1.05 (3H, d, $J$ = 7.5 Hz, H-10); $\delta_C$ (100 MHz, $CDCl_3$), 175.4 (C24), 170.9 (C8 or C26), 170.2 (C8 or C26), 72.5 (C3), 66.9 (C7), 57.8 (C12 or C13), 57.6 (C12 or C13), 51.3 (C25), 45.1 (C4), 29.8 (C14), 27.1 (C2), 21.4 (C9 or C27), 21.3 (C9 or C27), 14.8 (C10). Data obtained were consistent with those reported in literature.

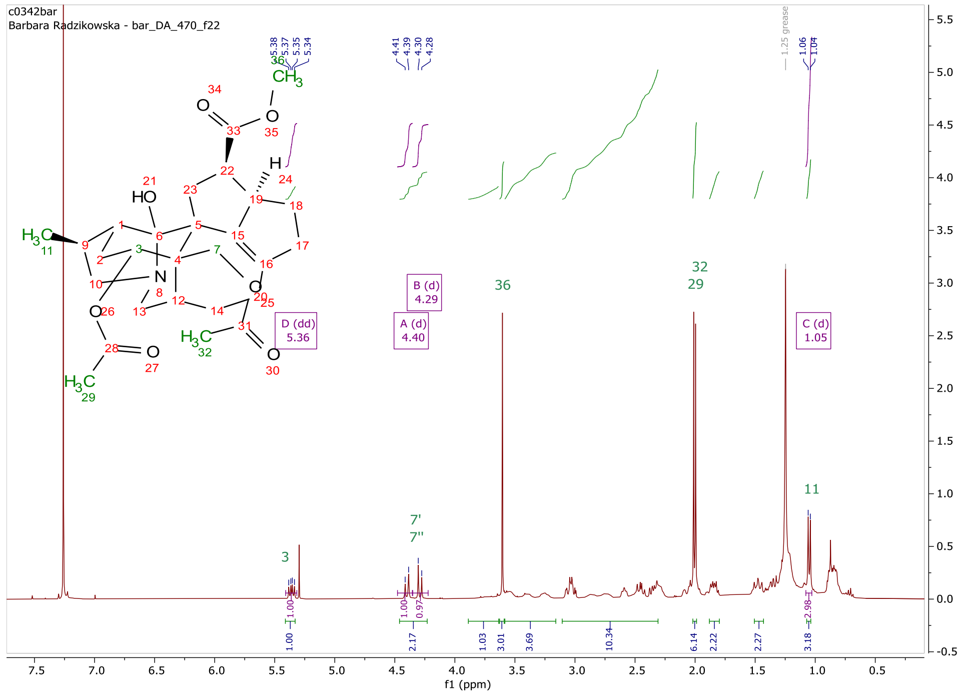


$^1$H NMR spectrum of yuzurimine.

#### Daphniphylline

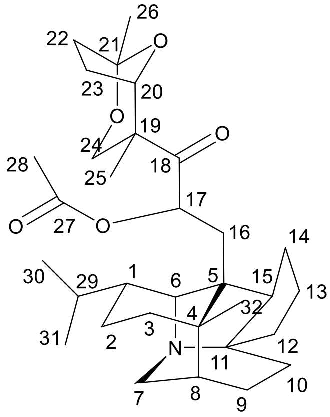

daphniphylline

$\delta_H$ (400 MHz, $CDCl_3$), 5.63 (1H, dd, *J* = 12.5. 3.0 Hz, H-17), 4.49 (1H, dd, *J* = 13.0, 2.0 Hz, H-24a), 3.72 (1H, d, *J* = 13.0 Hz, H-17b), 3.49-3.29 (1H, m, H-6), 3.24-2.92 (2H, m, H-7), 2.11 (3H, s, H-28), 1.40 (3H, s, H-26), 1.33 (3H, s, H-25 or H-32), 1.01 (3H, d, J = 7.0 Hz, H-30 or H-31), 0.99 (3H, d, J = 5.0 Hz, H-30 or H-31), 0.91 (3H, s, H-25 or H-32). Data obtained were consistent with those reported in literature.

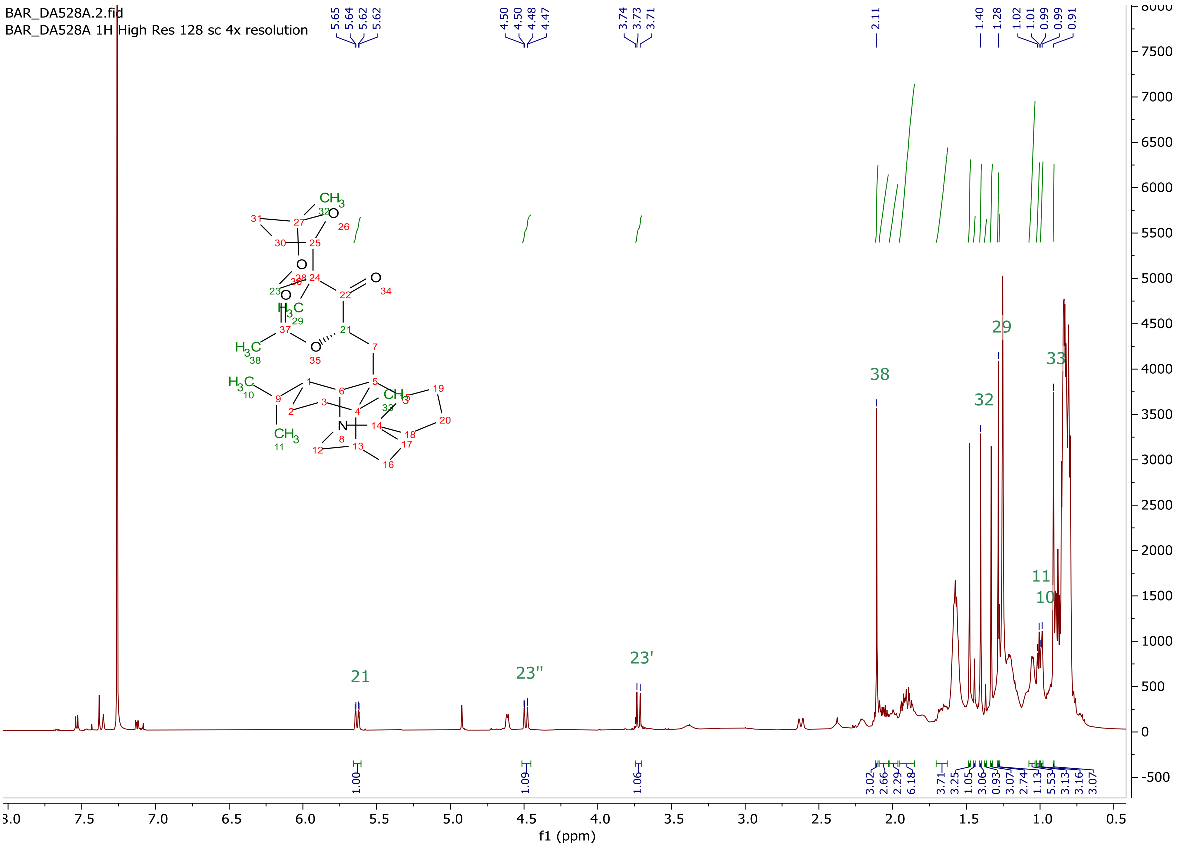


$^1$H NMR spectrum of daphniphylline with impurities.
