## Supplementary material for "Integrative metabolomics reveal the organisation of alkaloid biosynthesis in *Daphniphyllum macropodum*": Submission_bioRxiv_SI.pdf

[1]Centre for Novel Agricultural Products (CNAP), Department of Biology, University of York, York, UK; [2]Department of Chemistry, University of York, York, UK; [3]Department of Statistics, University of Warwick, Coventry, UK; [4]Department of Biology, University of York, York, UK; [5]Alan Turing Institute, London, UK; [6]Metabolomics and Proteomics Lab, Bioscience Technology Facility, Department of Biology, University of York, York, UK.

**Figures**

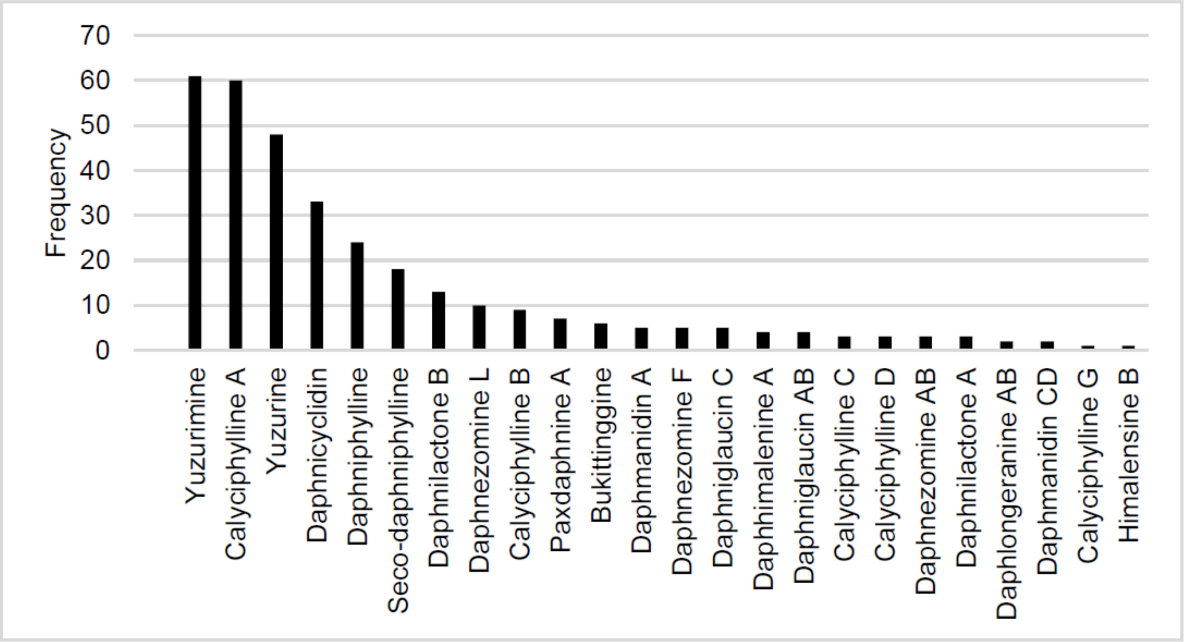


Supplementary Figure 1. Frequency of *Daphniphyllum* alkaloid compound subtype annotations in the *in silico* metabolite database collated from literature and databases.

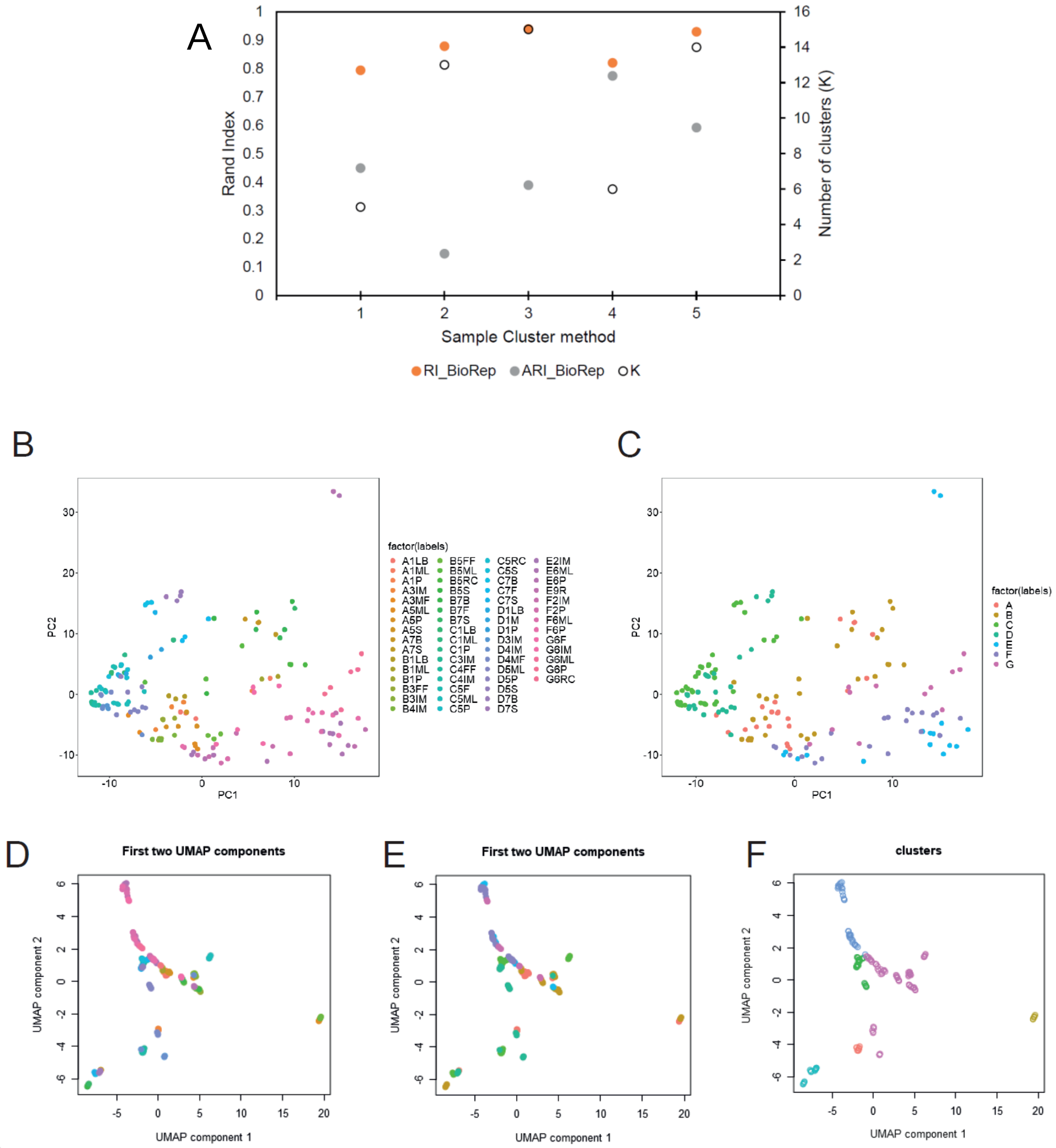


Supplementary Fig. 2. Sample clustering. A. Comparison of Rand index, Adjusted Rand Index and number of clusters (K) in a selected subset of clustering methods. For details of parameters see Supplementary Table 3. Sample Cluster method 4 was selected as it had the highest ARI. B and C. First two principle components of samples coloured by plant-tissue annotation (B) and plant (C). D, E and F. Projection of samples on first two UMAP components coloured by plant-tissue annotation (D), plant (E) and cluster assignment (F).

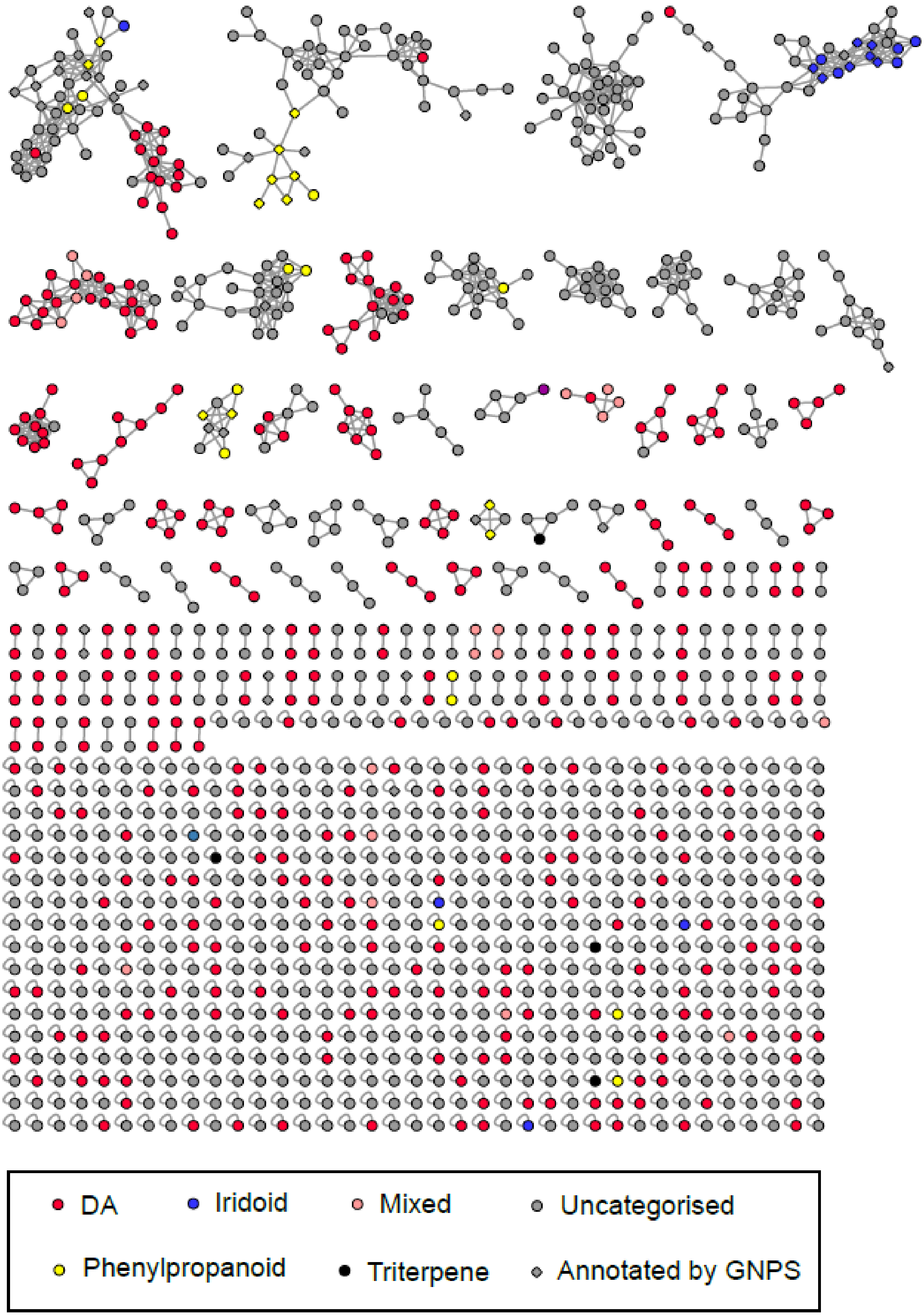


Supplementary Fig. 3. Feature based molecular network from GNPS. The network was constructed using composite HCD $MS^2$ data. Peak chemical type annotations are depicted by coloured nodes. Diamond shaped peak nodes represent peaks which have been annotated by matches to GNPS spectral libraries.

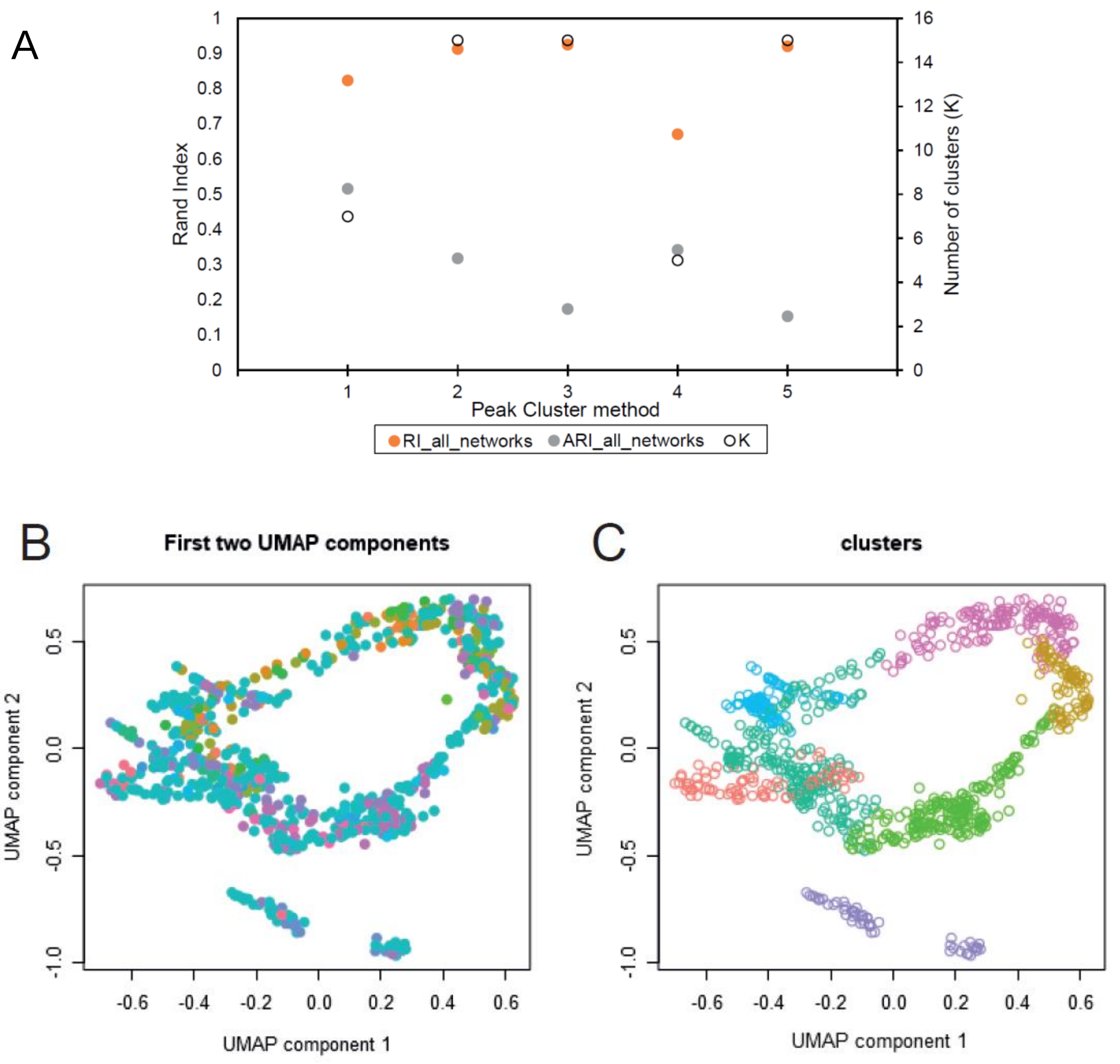


Supplementary Fig. 4. Peak clustering. A. Comparison of Rand index, Adjusted Rand Index and number of clusters (K) in a selected subset of clustering methods. For details of parameters see Supplementary Table 4. Peak cluster method 1 was selected as it had the highest ARI. B and C. Projection of peaks on first two UMAP components coloured by network-type annotation used for ARI calculation (B) and by cluster assignment (C).

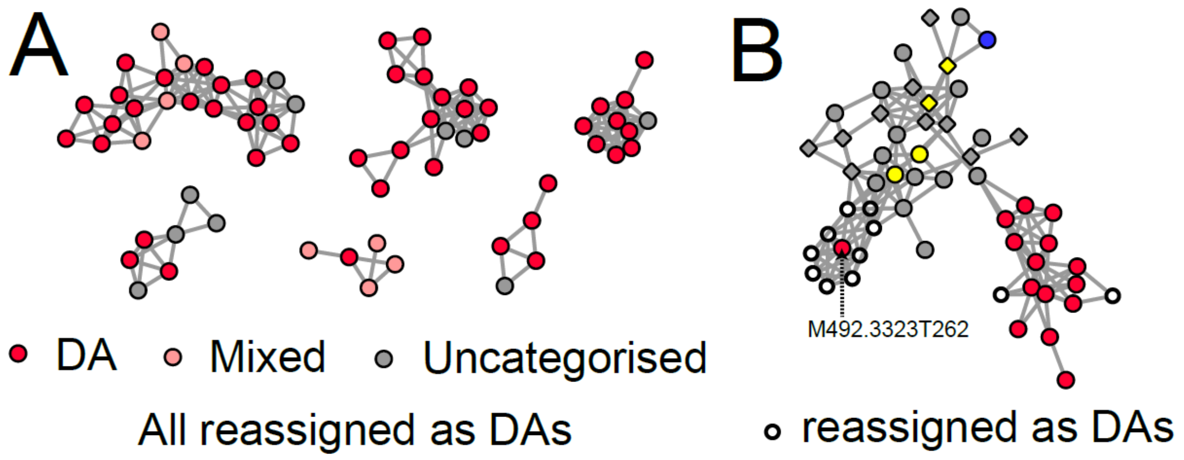


Supplementary Fig. 5. Peaks re-annotated as DAs due to position in molecular network. A. Selected subnetworks which contained DAs where all connected peaks were re-annotated as DAs. The annotation “mixed” indicated the peaks were annotated as multiple compounds based on MS1 library matches (often DA and a non-DA alkaloid). B. Complex subnetwork containing multiple compound types. The previously uncategorised peaks depicted with an unfilled circle were reannotated as DAs due to their proximity to DA peaks. Details of peaks connected to M492.3323T262 are described in Supplementary Fig. 6.

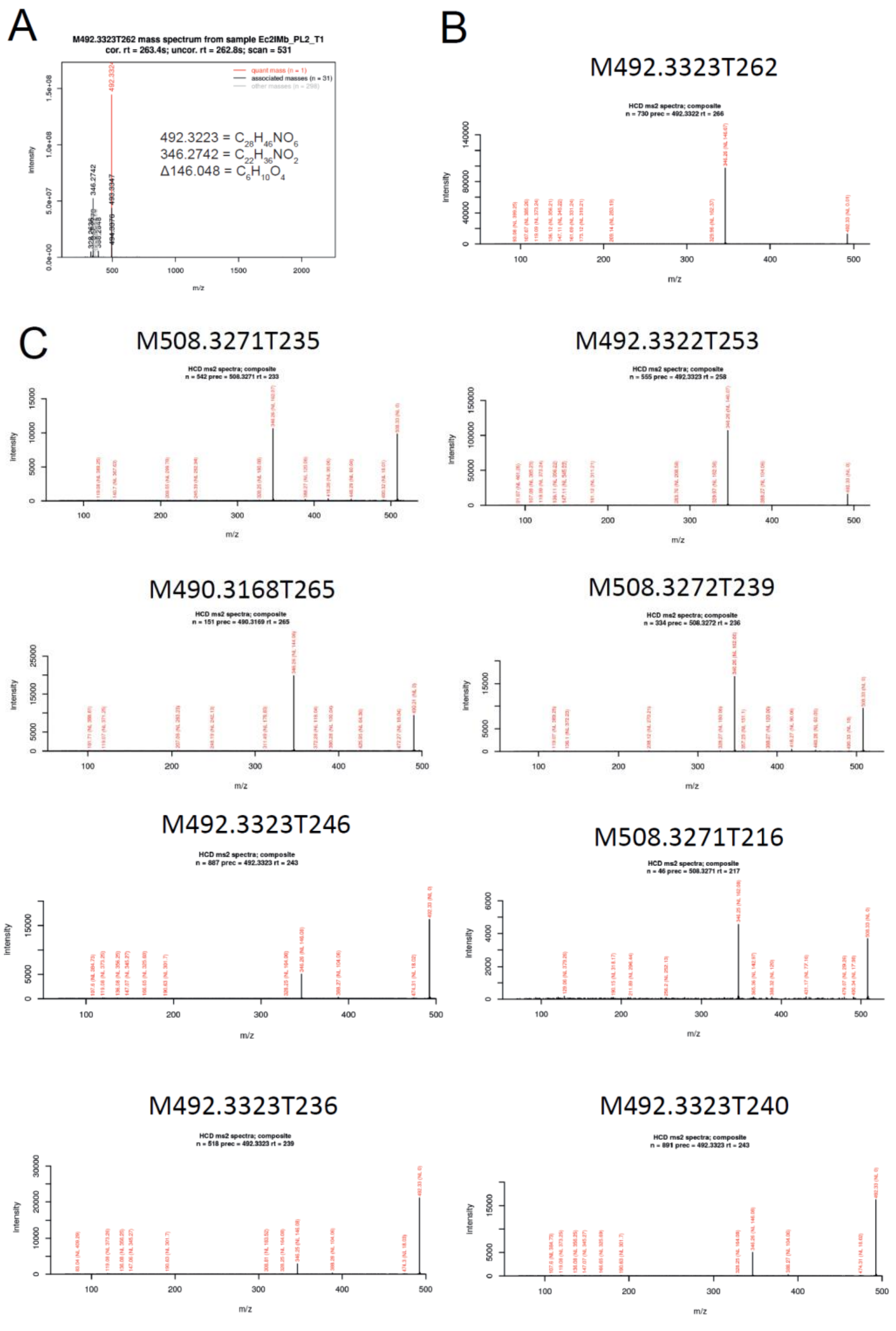


Supplementary Fig. 6. Assignment of M492.3323T262 and related peaks as DAs. A. $MS^1$ spectrum of M492.3323T262 showing in-source fragmentation to 346.2742. B. HCD composite $MS^2$ spectrum of M492.3323T262. C. HCD composite $MS^2$ spectra of peaks connected to M492.3323T262 on the GNPS network (see Supplementary Fig. 5) highlighting similar fragmentation behaviour.

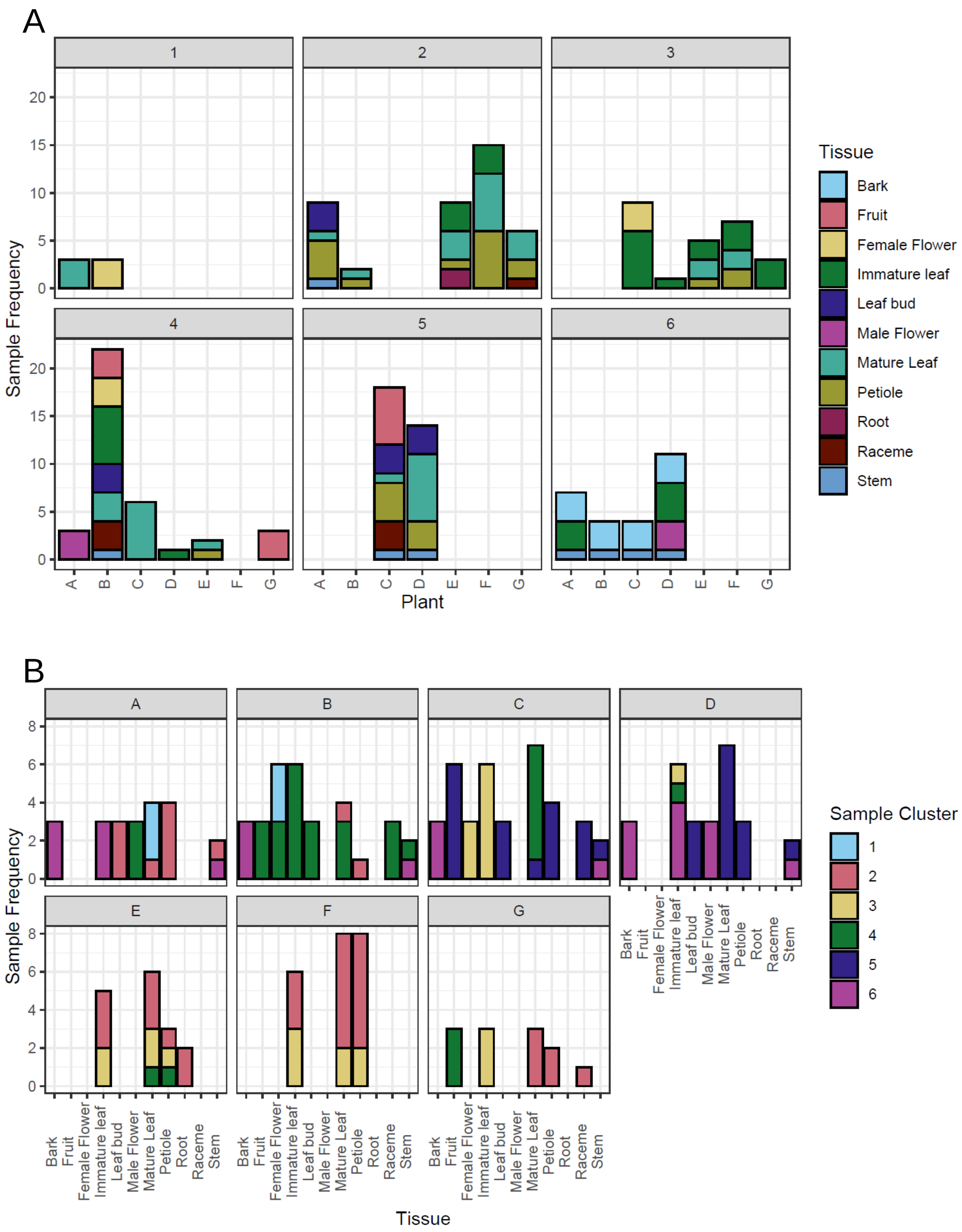


Supplementary Fig. 7. Content of the six sample clusters. A. Sample cluster composition, with plant origin (A-G) on the x-axis and tissue type coloured. B. Assignment of samples from plants (A-G), with tissue on the x-axis and Sample cluster assignment coloured.

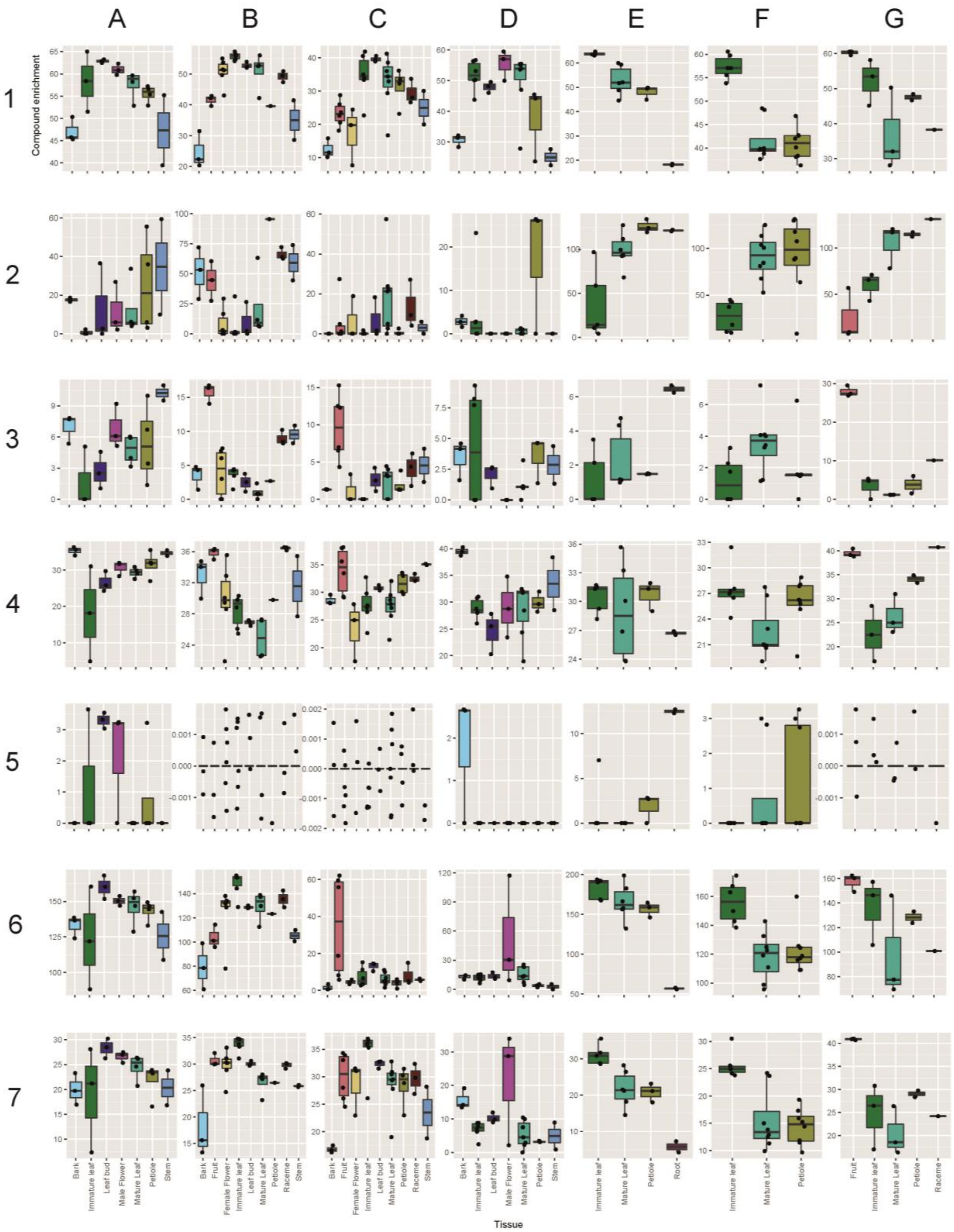


Supplementary Fig. 8. Enrichment of peak clusters across tissues within each plant. Plants grouped in columns and peak clusters in rows. Y-axis shows compound enrichment. All samples are plotted as points and the distribution as box-plots. Numbers of samples per plant-tissue combination varies. Kruskal-Wallis chi-squared tests and Dunn post-hoc tests can be found in Supplementary Data 4.

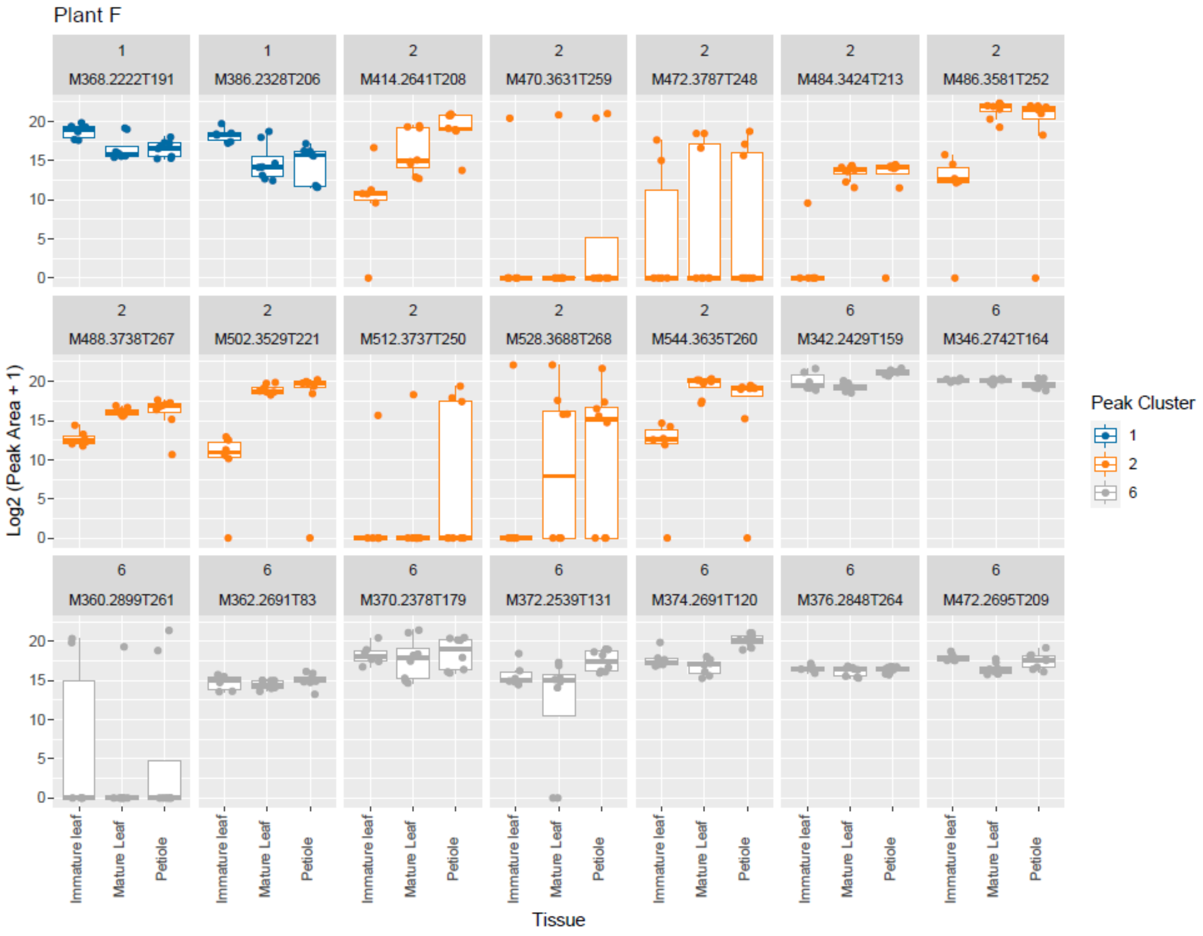


Supplementary Fig. 9. Distribution of most abundant DAs across plant F samples. These are the peaks from the LC-MS matching MALDI-MS ions (Supplementary Table 10). Note the different distribution of peaks from peak clusters across the tissue types, with peak cluster 1 highest in immature leaf and peak cluster 2 low in immature leaf. Y-axis is log2 transformed peak area.

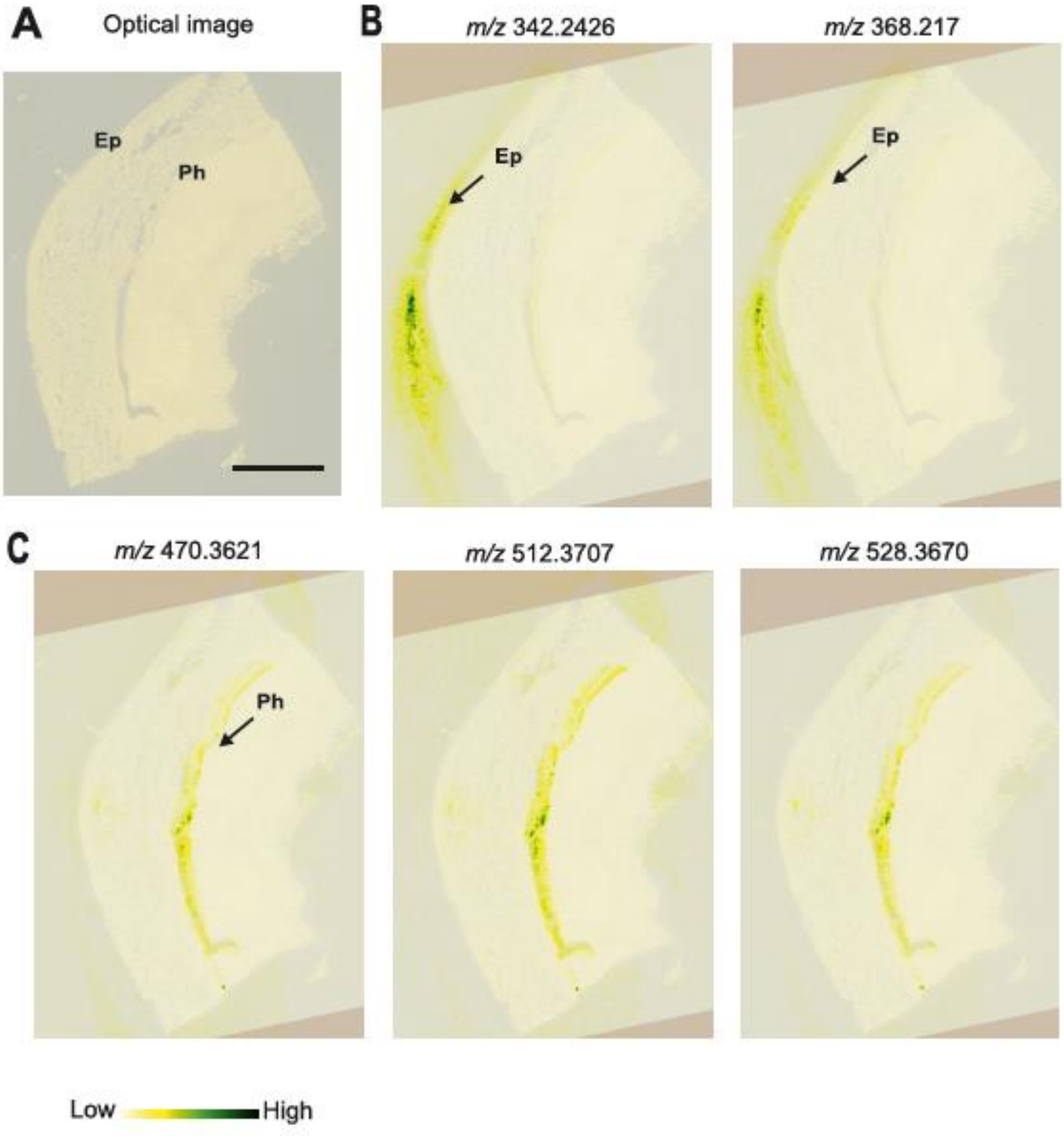


Supplementary Fig. 10. Ion intensity maps of selected alkaloids detected as $[M+H]^+$ ions with the MALDI-MS in the cross section of *D. macropodum* stem, colour bar represents MS signal intensity, positions of the epidermis (Ep) and the phloem region (Ph) are indicated. Panel A; optical image, Panel B; alkaloid presence in the epidermis, Panel C; alkaloid presence in the phloem region, size bar = 1000 µm. Data analysed in sensitivity mode with and laser energy 80.0 J $cm^{-2}$.

**Supplementary Tables**

Supplementary Table 1. Standards obtained for use in the LC-MS experiments.

| Compound name | Type | Chemical formula | Source |
|---|---|---|---|
| Daphylloside | Iridoid | $C_{19}H_{26}O_{12}$ | ChemFaces |
| Yuzurimine | DA | $C_{27}H_{37}NO_7$ | Purified in-house |
| Daphniphylline | DA | $C_{32}H_{49}NO_5$ | Purified in-house |
| Asperuloside | Iridoid | $C_{18}H_{22}O_{11}$ | Merck |
| Puerarin | Phenylpropanoid | $C_{21}H_{20}O_9$ | Merck |
| Luteolin | Phenylpropanoid | $C_{15}H_{10}O_6$ | Merck |
| Naringin | Phenylpropanoid | $C_{27}H_{32}O_{14}$ | Merck |
| Kaempferol | Phenylpropanoid | $C_{15}H_{10}O_6$ | Merck |
| Asperulosidic acid | Phenylpropanoid | $C_{18}H_{24}O_{12}$ | Generon |
| 5,7-dihydroxychromone | Phenylpropanoid | $C_9H_6O_4$ | Merck |
| Rutin | Phenylpropanoid | $C_{27}H_{30}O_{16}$ | Merck |
| Naringenin | Phenylpropanoid | $C_{15}H_{12}O_5$ | Merck |
| Quercetin | Phenylpropanoid | $C_{15}H_{10}O_7$ | Merck |
| Sarothrin | Phenylpropanoid | $C_{18}H_{16}O_8$ | Merck |
| Apigenin | Phenylpropanoid | $C_{15}H_{10}O_5$ | Merck |
| Kaempferol 3-rutinoside | Phenylpropanoid | $C_{27}H_{30}O_{15}$ | Generon |
| Deacetylasperulosidic acid | Iridoid | $C_{16}H_{22}O_{11}$ | Generon |

Supplementary Table 2. Plants used in this study.

| Code | species | var | Current Location | Supplier | Planted | Sex | Accession |
|---|---|---|---|---|---|---|---|
| A | *macropodum* | | Ray Wood, Castle Howard | Treseder Nurseries | 1978 | male | 197831591 |
| B | *macropodum* | *humile* | Ray Wood, Castle Howard | Jim Russell | 1989 | female | 19879448 |
| C | *macropodum* | | Ray Wood, Castle Howard | Jim Russell | 1989 | female | 20064193 |
| D | *macropodum* | | Ray Wood, Castle Howard | Jim Russell | 1989 | male | 20064194 |
| E | *macropodum* | | University of York | Burncoose nurseries | 2018 | not flowering | NA |
| F | *macropodum* | | University of York | Architectural plants | 2018 | not flowering | NA |
| G | *macropodum* | *humile* | University of York | Crûg Farm Plants | 2018 | female | BSWJ11232 |

| Code | Origin | Comments |
|---|---|---|
| A | China/Japan/Korea/Taiwan | Horticultural origin, exact geographic origin unknown |
| B | Japan | |
| C | Fanjin Shan, Guizhou, China | |
| D | Fanjin Shan, Guizhou, China | |
| E | China/Japan/Korea/Taiwan | Horticultural origin, exact geographic origin unknown, seed taken from mature tree on site in UK |
| F | China/Japan/Korea/Taiwan | Horticultural origin, exact geographic origin unknown, seed taken from mature tree on site in UK |
| G | B&SWJ 2005 5th expedition, Japan | |

Supplementary Table 3. Selected sample clustering parameters, showing representative examples with highest ARI (extracted from list of 7529).

| No | usePCA? | numPCs | useUMAP? | numNeighbours | numadjsComponents | clusteringAlgorithm | K | RI_BioRep | ARI_BioRep |
|---|---|---|---|---|---|---|---|---|---|
| 1 | FALSE | 0 | TRUE | 11 | 16 | hierarchical_complete | 5 | 0.79380997 | 0.449445911 |
| 2 | FALSE | 0 | TRUE | 15 | 16 | hierarchical_median | 13 | 0.87814732 | 0.1476109 |
| 3 | TRUE | 5 | TRUE | 5 | 2 | hierarchical_complete | 15 | 0.939254022 | 0.389390945 |
| 4 | TRUE | 10 | TRUE | 5 | 4 | K-means | 6 | 0.819565688 | 0.773953662 |
| 5 | TRUE | 7 | TRUE | 11 | 4 | hierarchical_complete | 14 | 0.92944232 | 0.591597359 |

Supplementary Table 4. Selected peak clustering parameters, showing representative examples with highest ARI (extracted from list of 7257)

| No | usePCA? | numPCs | useUMAP? | numNeighbours | numadjsComponents | clusteringAlgorithm | K | RI | ARI |
|---|---|---|---|---|---|---|---|---|---|
| 1 | FALSE | 0 | TRUE | 17 | 12 | hierarchical_complete | 7 | 0.823915266 | 0.516075408 |
| 2 | TRUE | 10 | TRUE | 15 | 2 | K-means | 15 | 0.912509713 | 0.318598511 |
| 3 | TRUE | 5 | TRUE | 13 | 2 | K-means | 15 | 0.924949904 | 0.174198009 |
| 4 | TRUE | 6 | TRUE | 17 | 2 | hierarchical_median | 5 | 0.671091482 | 0.342790796 |
| 5 | TRUE | 5 | TRUE | 9 | 2 | hierarchical_complete | 15 | 0.920394226 | 0.153558583 |

Supplementary Table 5. LC-MS peaks corresponding to standards or $MS^2$ fragmentation patterns. All $MS^2$ matches are from M.-M. Cao et al. *Curr. Pharm. Anal.* 12, 249–257 (2016).

| Peak | Compound | Type | Subtype | Origin | Comments |
|---|---|---|---|---|---|
| M303.0500T57 | Rutin | Phenylpropanoid | Flavonoid/lignan | Standard | |
| M595.1661T60 | Kaempferol-3-rutinoside | Phenylpropanoid | Flavonoid/lignan | Standard | |
| M287.0551T60 | Luteolin | Phenylpropanoid | Flavonoid/lignan | Standard | Main peak |
| M287.0551T56 | Luteolin | Phenylpropanoid | Flavonoid/lignan | Standard | Peak shoulder |
| M417.1182T57 | Puerarin C | Phenylpropanoid | Flavonoid/lignan | Standard | Main peak |
| M417.1181T60 | Puerarin C | Phenylpropanoid | Flavonoid/lignan | Standard | Peak shoulder |
| M450.1608T57 | Asperulosidic acid | Iridoid | Iridoid | Standard | |
| M287.0551T67 | Kaemferol | Phenylpropanoid | Flavonoid/lignan | Standard | |
| M273.0758T78 | Naringin | Phenylpropanoid | Flavonoid/lignan | Standard | |
| M464.1765T75 | Daphylloside | Iridoid | Iridoid | Standard | |
| M486.2489T203 | Daphtenidine D-like | DA | Yuzurimine | MS2 | Tentative match |
| M486.2489T157 | Daphtenidine D-like | DA | Yuzurimine | MS2 | Tentative match |
| M470.2539T211 | Yuzurimine | DA | Yuzurimine | Standard | In source loss of water |
| M384.2172T236 | Daphhimalenine B-like | DA | Yuzurimine | MS2 | Tentative match |
| M384.2172T240 | Daphhimalenine B-like | DA | Yuzurimine | MS2 | Tentative match |
| M384.2172T268 | Daphhimalenine B-like | DA | Yuzurimine | MS2 | Tentative match |
| M432.2747T267 | Daphmacromine B-like | DA | Yuzurine | MS2 | Tentative match |
| M400.2120T162 | Daphnezomine U | DA | Daphnezomine F | MS2 | Tentative match |
| M404.2433T140 | Daphnezomine K-like | DA | Yuzurimine | MS2 | Tentative match |
| M472.2695T209 | Deoxyyuzurimine | DA | Yuzurimine | MS2 | Good match |
| M382.2018T173 | Yuzurimine C-like | DA | Yuzurimine | MS2 | Tentative match |
| M382.2015T182 | Yuzurimine C-like | DA | Yuzurimine | MS2 | Tentative match |
| M418.2590T235 | Daphmacromine E-like | DA | Yuzurine | MS2 | Tentative match |
| M418.2590T239 | Daphmacromine E-like | DA | Yuzurine | MS2 | Tentative match |
| M418.2591T243 | Daphmacromine E-like | DA | Yuzurine | MS2 | Tentative match |

Supplementary Table 6. Kruskal–Wallis tests (non-parametric ANOVA) detecting significant differences in peak cluster enrichment across plants and sample clusters. Dunn post-hoc tests are shown in Supplementary Data 3 and the resulting pairwise significance bins in Supplementary Table 7 and 8.

| Peak cluster | Plant variation, df = 6 | | Sample cluster variation, df = 5 | |
|---|---|---|---|---|
| | $X^2$ | p-value | $X^2$ | p-value |
| 1 | 26.175 | 0.0002066 | 34.636 | 1.778e-06 |
| 2 | 92.723 | <2.2e-16 | 64.962 | 1.141e-12 |
| 3 | 13.239 | 0.0394 | 23.608 | 0.0002581 |
| 4 | 36.006 | 2.749e-06 | 41.541 | 7.293e-08 |
| 5 | 15.525 | 0.01654 | 14.273 | 0.01397 |
| 6 | 126.34 | <2.2e-16 | 65.339 | 9.532e-13 |
| 7 | 125.18 | <2.2e-16 | 28.171 | 3.37e-05 |

Supplementary Table 7. Pairwise significance bins for peak cluster and plant compound enrichment. Groups sharing letters do not have significantly different means. Determined from Dunn post-hoc test with Benjamini-Hochberg multiple test correction. Boxplot is Fig 4A, Kruskal–Wallis test Supplementary Table 6 and full details of the Dunn post-hoc test in Supplementary Data 3.

| Plant | Peak cluster | | | | | | |
|---|---|---|---|---|---|---|---|
| | 1 | 2 | 3 | 4 | 5 | 6 | 7 |
| A | ab | a | abcd | acdef | a | abd | ae |
| B | abd | a | acd | bd | b | ab | bf |
| C | ab | b | bc | acd | b | c | cf |
| D | abcd | b | abc | abcd | b | c | de |
| E | cd | c | abcd | aef | ab | ad | ae |
| F | cd | c | abcd | af | ab | abd | ade |
| G | bcd | c | ad | abcdf | ab | abd | bcf |

Supplementary Table 8. Pairwise significance bins for peak cluster and sample cluster compound enrichment. Groups sharing letters do not have significantly different means. Determined from Dunn post-hoc test with Benjamini-Hochberg multiple test correction. Boxplot is Fig 4B, Kruskal–Wallis test Supplementary Table 6 and full details of the Dunn post-hoc test in Supplementary Data 3.

| Sample cluster | Peak cluster | | | | | | |
|---|---|---|---|---|---|---|---|
| | 1 | 2 | 3 | 4 | 5 | 6 | 7 |
| 1 | ab | acdef | ab | ab | abcd | ab | ab |
| 2 | abc | b | ab | a | ad | ab | ab |
| 3 | abc | acdf | b | a | bcd | abd | b |
| 4 | abc | acde | a | b | abcd | ab | b |
| 5 | bc | aef | a | b | bc | cd | ab |
| 6 | d | acef | a | b | abd | bcd | a |

Supplementary Table 9. Kruskal–Wallis tests (non-parametric ANOVA) detecting significant differences in peak cluster enrichment across tissues, with data from plants ABEFG combined. Dunn post-hoc tests are shown in Supplementary Data 4.

| Peak cluster | Tissue variation, df = 10 | |
|---|---|---|
| | $X^2$ | p-value |
| 1 | 49.288 | 3.606e-07 |
| 2 | 45.875 | 1.511e-06 |
| 3 | 48.492 | 5.047e-07 |
| 4 | 48.113 | 5.918e-07 |
| 5 | 34.22, | 0.0001695 |
| 6 | 27.813 | 0.001934 |
| 7 | 55.324 | 2.747e-08 |

Supplementary Table 10. Representative DAs detected in *D. macropodum* tissues using MALDI-MS and the corresponding LC-MS peaks and peak clusters. See Supplementary Fig. 9 for LC-MS peak data across plant F samples.

| MALDI m/z | Formula | Theoretical m/z $[M+H]^+$ | Δppm | Matching LCMS peak | Peak cluster |
|---|---|---|---|---|---|
| 342.2426 | C22H31NO2 | 342.2433 | -2.051172 | M342.2429T159 | 6 |
| 346.2730 | C22H35NO2 | 346.2746 | -4.62061 | M346.2742T164 | 6 |
| 360.2573 | C22H33NO3 | 360.2539 | 9.437788 | M360.2899T261 | 6 |
| 362.2674 | C22H35NO3 | 362.2695 | -5.796789 | M362.2691T83 | 6 |
| 368.217 | C23H29NO3 | 368.2226 | -15.208192 | M368.2222T191 | 1 |
| 370.2333 | C23H31NO3 | 370.2382 | -13.234723 | M370.2378T179 | 6 |
| 372.2513 | C23H33NO3 | 372.2539 | -6.98448 | M372.2539T131 | 6 |
| 374.2678 | C23H35NO3 | 374.2695 | -4.584931 | M374.2691T120 | 6 |
| 376.2856 | C23H37NO3 | 376.2852 | 1.063023 | M376.2848T264 | 6 |
| 386.2364 | C23H31NO4 | 386.2331 | 8.544063 | M386.2328T206 | 1 |
| 414.2611 | C25H35NO4 | 414.2644 | -8.038344 | M414.2641T208 | 2 |
| 470.3621 | C30H47NO3 | 470.3634 | -2.76382 | M470.3631T259 | 2 |
| 472.2658 | C27H37NO6 | 472.2699 | -8.681476 | M472.2695T209 | 6 |
| 472.3722 | C30H49NO3 | 472.3791 | -14.606912 | M472.3787T248 | 2 |
| 484.3440 | C30H45NO4 | 484.3426 | 2.725343 | M484.3424T213 | 2 |
| 486.3571 | C30H47NO4 | 486.3583 | -2.467317 | M486.3581T252 | 2 |
| 488.3658 | C30H49NO4 | 488.3740 | -16.790411 | M488.3738T267 | 2 |
| 502.3462 | C30H47NO5 | 502.3532 | -14.022006 | M502.3529T221 | 2 |
| 512.3707 | C32H49NO4 | 512.3740 | -6.440608 | M512.3737T250 | 2 |
| 528.3670 | C32H49NO5 | 528.3689 | -3.595972 | M528.3688T268 | 2 |
| 544.3588 | C32H49NO6 | 544.3638 | -9.185034 | M544.3635T260 | 2 |

**Supplementary Data**

Supplementary Data 1: Sample description and metadata

Supplementary Data 2: R-script for clustering algorithm.

Supplementary Data 3: Dunn post-hoc tests for peak cluster enrichment across plants and sample clusters.

Supplementary Data 4: Kruskal–Wallis test and Dunn post-hoc tests for peak cluster enrichment across tissues.

Supplementary Data 5: NMR analysis of isolated alkaloids.
